# Tuning the performance of a TphR-based terephthalate biosensor with a design of experiments approach

**DOI:** 10.1101/2024.06.26.600737

**Authors:** Guadalupe Alvarez Gonzalez, Micaela Chacón, Thomas Butterfield, Neil Dixon

## Abstract

Transcription factor-based biosensors are genetic tools that aim to predictability link the presence of a specific input stimuli to a tailored gene expression output. The performance characteristics of a biosensor fundamentally determines its potential applications. However, current methods to engineer and optimise tailored biosensor responses are highly nonintuitive, and struggle to investigate multidimensional sequence/design space efficiently. In this study we employ a design of experiments (DoE) approach to build a framework for efficiently engineering activator-based biosensors with tailored performances, and we apply the framework for the development of biosensors for the polyethylene terephthalate (PET) plastic degradation monomer terephthalate (TPA). We simultaneously engineer the core promoter and operator regions of the responsive promoter, and by employing a dual refactoring approach, we were able to explore an enhanced biosensor design space and assign their causative performance effects. The approach employed here serves as a foundational framework for engineering transcriptional biosensors and enabled development of tailored biosensors with enhanced dynamic range and diverse signal output, sensitivity, and steepness. We further demonstrate its applicability on the development of tailored biosensors for primary screening of PET hydrolases and enzyme condition screening, demonstrating the potential of statistical modelling in optimizing biosensors for tailored industrial and environmental applications.

**Graphical Abstract**. Employment of a DoE framework for fine-tuning biosensor performance.

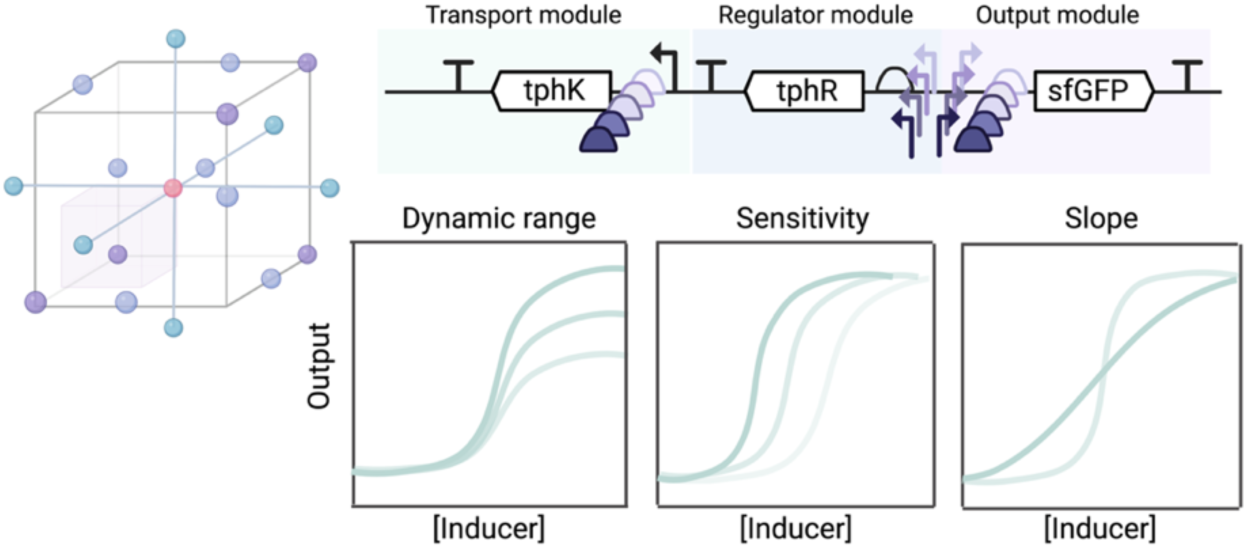

**Highlights:**

- Bioinformatic mining of allosteric transcription factors to produce TPA biosensors
- Efficient sampling of complex sequence-function relationships of genetic circuits
- Modelling to learn and optimise biosensor genetic circuits
- Application of biosensors for primary and secondary enzyme screening applications

## 1. Introduction

Genetically encoded biosensors are widely used regulatory tools that can enable precise measurement and control of biological behaviour. These inducible systems respond to specific input stimuli by regulating the expression of an output gene(s), such as a reporter protein or catabolic pathway, in accordance with the available input concentration. Given the large variety of allosteric transcription factors (aTF) found in nature and the diverse range of stimuli and effector metabolites that they can recognize, transcriptional biosensors are widely adopted as a valuable class of regulatory tool ^1–4^. Depending on the relationship between the inducer ligand, the aTF and DNA operator site, different mechanisms of transcriptional control exist, including simple repression-de-repression and activator systems ^5,6^. Activator aTFs generally act by binding to the operator region co-located a corresponding promoter sequence, and upon ligand binding, recruit RNA polymerase (RNAP) through direct interactions with the RNAP sigma factor, permitting subsequent transcription initiation ^6–8^. When exposed to varying concentration of ligand, the sigmoidal gene expression response curve elicited by aTF-dependent biosensors can be described by simple mathematical models, like the Hill function. These models can help define biosensor performance features including the minimal and maximal response, the half maximal effective concentration (EC_50_) and the Hill coefficient (n_H_), which in turn define the biosensor dynamic range (max/min response), sensitivity and curve steepness, respectively ^5,9^. The potential application of a biosensor is therefore dictated by the performance of its dose-response curve. For instance, biosensors displaying a high dynamic range can allow confident discernment of variants in high-throughput enzyme/strain screening applications, or serve as regulatory components for protein production systems upon specific substrate induction^10–12^. Moreover, biosensors that exhibit greater sensitivity and a steep (digital) response curve could be used as primary screening tools, while those with a less steep (analogue) response curved could provide specific and graded output responses ideal for secondary screening between closely related enzyme variants or distinct conditions^4,13^. Other custom dose-response curves could find further numerous modern applications, such as *in situ* diagnosis or monitoring, conditional control of cell differentiation and synthetic cell–cell communication devices^2,6,14^. Whatever the application, biosensors are required to be fully tuneable and modularizable, so that their individual genetic components can be carefully attenuated to achieve the desired performance parameters.

In general, after the initial biosensor construction, iterative rounds of adjustment are often necessary to optimise the expression levels and stoichiometry of individual regulatory component in order to achieve the required performance. Most holistic design and optimization efforts to date have relied upon iterative engineering of said individual components, typically through protein or promoter engineering, or via mathematical or mechanistic modelling ^5,15–17^. Achieving predictable and reliable performance in this way thus requires a deep understanding of the intricate interplay between the specific biosensor’s components, as changes to one element can simultaneously affect all responses.

To address such multi-factorial biological engineering challenges, an expanding body of research has turned to the use of employing structured multivariate and statistical experimentation to drive data-led design, characterization, and optimization of biological systems^18–24^. Design of experiments (DoE) is one such powerful statistical methodology that can be applied to guide the design and optimization of bioprocess conditions or desired genetic constructs^19,25^. It employs statistical modelling to systematically analyse a multifactorial experimental design space with defined variables, while suggesting a minimal number of experiments needed to study the impact of these variable upon overall system performance. More crucially, DoE can achieve significant optimisation and understanding of a system without requiring extensive *a priori* knowledge of the precise mechanisms that underlie it. Therefore, various DoE approaches have been successfully applied to guide and optimise the efficiency of metabolic engineering and bioprocess development design^19,24,26^. We previously implemented this methodology to optimise the dynamic range performance of a simple biosensor operating via a repression-de-repression mechanism, by varying the expression levels of the regulator (repressor) and the output gene (*eGFP*) ^18^. However, more complex regulatory systems, such as activator-based biosensors can display strong interdependency between their regulatory components, where the function of an individual component might rely directly on a second component, or more. Given the considerable effort required to build new transcriptional biosensors and their unpredictable performance, a comprehensive framework that enables the exploration of how each component influences each response could streamline the overall design process, minimising the time and effort needed to determine the system configuration leading to a required optimal performance.

Among emerging applications of genetically encoded biosensors, these tools are expected to play a significant role in advancing and supporting the biological synthesis and degradation of plastics, contributing to achieving greater waste circularity ^27–29^. For example, to support the transition away from petrochemical-based to renewable feedstocks, biosensors have been applied to facilitate metabolic engineering efforts to produce drop-in monomer precursors and/or for bio-based plastic production ^4,30,31^. Additionally, biosensors can be highly powerful screening tools to discovery and optimize novel enzymes for the efficient breakdown of plastics^27,28^. Here, we applied DoE as a design framework to achieve customised performance of activator-based biosensors. Specifically, we harness this workflow to build biosensors for terephthalic acid (TPA) with custom performances in *Pseudomonas putida* KT2440. TPA is an aromatic building block of polyethylene terephthalate (PET) plastic, and is released after enzymatic or chemical deconstruction of PET^32–34^. Therefore, optimal TPA detection methods find immediate application in the discovery and engineering of novel PET hydrolases, or the dynamic regulation of biosynthetic pathways towards PET valorisation^35–40^. Here, we identified and characterized novel regulatory components to build a fully modularized TPA biosensor design, which we then used to build an efficient TPA-responsive promoter library that covers a wide range of sensing strengths, dynamic ranges, and sensitivities. Using a dual promoter library ranking approach, we then used regression modelling to identify the main factors affecting the simultaneous performance responses of the resulting biosensors. Collectively, the dual approach employed here serves as a framework for activator-based biosensors and enabled development of tailored biosensors with enhanced dynamic range and diverse signal output, sensitivity, and steepness. Finally, two biosensors with distinct digital and analogue performance responses were used in both i) a primary screening application to triage a library of hydrolases for PET construction, and ii) a subsequent enzyme condition screening application.

## 2. Methods

### 2.1 Materials and strains

*Escherichia coli* DH5α (NEB) was used for cloning and DNA assembly of all mutant variants and constructs. *Pseudomonas putida* KT2440 was used for characterization of all parts and constructs and for all modelling experiments. All strains were grown in liquid Luria−Bertani (LB) media or agar plates supplemented with either 100 μg/ml (*E. coli*) or 500 μg/ml (*P. putida*) carbenicillin and incubated overnight at 37°C or 30°C, respectively. Disodium terephthalic acid (Thermo Scientific) stock solutions were prepared in sterile water. Esterases from *Clostridium thermocellum* and *Cellvibrio japonicus* were purchased from Prozomix Limited, lipases from *Pseudomonas fluorescens*, *Aspergillus niger*, *Candida rigosa*, *Mucor miehei*, *Burkholderia cepacia*, *Rhizopus niveus*, *Candida antarctica* (CALB), and porcine liver were purchased from Merck. The LCC-ICCG cutinase was produced and purified as previously described ^40^.

### 2.2 Molecular cloning

The sequences of all plasmids and oligonucleotides used in this study can be found in Table S4 and Table S5, respectively. All DNA and oligonucleotide synthesis was performed by Integrated DNA Technologies (IDT). DNA amplification for cloning was performed using Q5 polymerase (NEB) and DNA amplification for colony PCR was performed using Phire II Green (Thermo Fisher). The pSEVA131 plasmid was used as the backbone for the generation of all biosensor constructs, and plasmid assembly was performed using NEBuilder HiFi DNA Assembly Mix (NEB), followed by transformation into chemically competent *E. coli* DH5α cells. Sanger sequencing (Eurofins Genomic, Germany) was performed for validation of correct construct assembly. For all characterization experiments, plasmids were transformed into *P. putida* by electroporation.

### 2.3 Phylogenetic analysis and promoter region motif identification

As previously reported ^41^, we found several putative tph operon homologues (*tph*-like operons) using BLAST-P to find homologues of the Comamonas sp. E6 (E6) *tphC* and then performing genomic context analysis to find operons enriched in other potential *tph* genes in addition to *tphC*. In most (17/22, 77%), of these *tph*-like operons, the *tphR* gene was located directly adjacent to another potential *tph* gene homologue with both genes being transcribed in opposite directions from a common intergenic region. In these cases, we considered the entire intergenic region between the *tphR* and the other *tph* gene to be a potential promoter region. These regions, together with that of E6, were searched for motifs using MEME ^42^ (classic mode, standard alphabet and ‘ANR’ motif distribution). The search was restricted to the sense strand and five motifs were selected for finding. A significant resulting motif was selected that overlapped the previously identified experimental motif ^43,44^ but generalised across the whole set of putative tph operons.

### 2.4 Part library validation, cloning and screening

The previously developed P_proB_ promoter and G10 RBS libraries^18^, were purified and transformed into *P. putida*. Overnight cultures of *P. putida* transformants were sub-cultured to an OD of 0.05 into 10 ml LB + carbenicillin before being aliquoted into 96 deep-well plates (DWPs) to a final volume of 500 µl. The DWPs were incubated at 30 °C, 1000 rpm, and 75 % humidity in a shaker incubator for 6 and 8 h. To measure optical density and fluorescence, culture pellets were washed and re-suspended in 500 µl PBS before being transferred to black clear bottom microtiter plates (BMTPs; Greiner). All density and fluorescence readouts were performed in a ClarioStar microplate reader (BMG). OD was measured at 700 nm, GFP was measured at λEx/λEm = 488/520 nm, and mCherry fluorescence was measured at λEx/λEm = 570/620 nm. Fluorescence was normalized to OD. Initial screening of the potential TPA sensitive promoter regions was performed similarly. Overnight cultures of *P. putida* transformants were sub-cultured into 10 ml LB + carbenicillin before being aliquoted into 96 deep-well plates (DWPs) to a final volume of 500 µl, with each well containing the relevant amount of TPA. The DWPs were incubated at 30 °C, 1000 rpm, and 75 % humidity in a shaker incubator for 16 hours. To measure optical density and fluorescence, culture pellets were washed and re-suspended in 500 µl PBS before being transferred to black clear bottom microtiter plates and quantified as outlined above.

The P_out_ promoter library was generated by linearisation of the original pTB4 plasmid and insertion of the GA229 degenerate single-stranded DNA oligo by isothermal assembly (Table S4). The resulting reaction mixture was transformed into *E. coli* DH5α, plated onto LB agar + carbenicillin, and incubated at 37 °C. A total of ∼7×10^4^ individual colonies were then pooled and the plasmid library DNA was isolated by midi-prep kit (Qiagen, UK). 10 μg of the resulting library DNA was then transformed into *P. putida* by electroporation, plated onto LB agar + carbenicillin, and incubated at 30 °C. Thirty colonies were individually sequenced for library quality control. A Colony Picker Qpix 400 (Molecular Devices, UK) was used to pick 5,000 individual colonies, and these were subsequently arrayed into 96 microtiter plates (MTPs; Greiner, UK) containing 200 μl of LB + carbenicillin per well. The MTPs were incubated for 16 h at 30 °C, 850 rpm, and 75 % humidity in a plate shaker incubator (Infors HT). Using a Hamilton robotic platform, 130 μl of the overnight library clones were transferred to a microtiter plate containing 130 μl of 50 % glycerol. These plates were briefly mixed in the plate-shaker (850 rpm, 5 min) and then stored at −80 °C for cryopreservation.

Screening of the resulting library clones was performed in two stages. First, stock MTP plates were thawed for 30 min., and using the Hamilton Robotic platform, 5μl were transferred into 96 DWPs containing 495 μl LB + carbenicillin. Plates were then grown for 16 h at 30°C, 850 rpm, and 75% humidity. 5 μl of culture from each well was then subcultured into four DWPs (total of 20 μl of culture from each well) containing LB + carbenicillin and either 0, 1, 25 or 1000 μM of terephthalic acid. These plates were grown for 16 h at 30°C, 850 rpm, and 75% humidity in a plate-shaker incubator. A total of 200 plates were then spun down, washed with PBS and finally re-suspended to 500 μl of PBS. 200 μl of the resulting resuspension were transferred to black clear bottom microtiter plates (BMTPs; Greiner) for optical density and fluorescence readout. This initial screening round was used to select a total of 226 relative functional clones (same or above functionality to original biosensor). In the next stage, this screening process was repeated where each clone variant was subcultured in triplicate. The resulting 226 variant library was similarly stored in MTPs for cryopreservation at −80°C and re-characterized for optical density and fluorescence.

### 2.5 DoE trial cloning and characterization by biosensor dose-response

Constructs conforming to the corresponding DoE Definitive Screening Designs (Table S1) were generated in three stages. First, the G10 RBS upstream of *tphK* was set to all relative library levels (RBS_trans_ -1, 0 and +1). The three resulting plasmids were linearised, removing the existing P_reg_ and P_out_/RBS_out_ sequences. Next, six overlapping fragments corresponding to all P_reg_ (P_proB_ -1, 0 and +1) and RBS_out_ levels (G10 -1, 0 and +1) were generated by PCR, and the final corresponding designs – except for P_out_ – were generated by inserting the desired genetic parts into the corresponding RBS_trans_ backbone. Finally, five overlapping P_out_ fragments (DR_out_ -1, 0 and +1 and EC -1 and +1) were synthesised, and these were subsequently integrated into the required linearized backbone according to the final DSD construct design. The final design plasmids were transformed into *P. putida* and streaked on LB plus carbenicillin. Three individual colonies from each construct were inoculated into 10 ml of LB plus carbenicillin and incubated overnight at 30 °C. The overnight cultures were diluted 1/100 into fresh LB and 450 μl were directly added into 96 DWPs containing 50 μl of TPA solution. The final concentrations for the titration were 0, 0.25, 0.5, 1, 5, 10, 25, 50, 100, 250 and 500 μM TPA. Plates were incubated for 16 h at 30 °C, 850 rpm, and 75% humidity. The DWPs were centrifuged, and pellets were washed twice before being resuspended in PBS. 100 μL of cell suspension was transferred to flat black MTP containing 100 μL of PBS, and GFP fluorescence and OD_600_ were measured as described above.

### 2.6 Data processing and modelling

Relative constitutive promoter (P_proB_) or RBS (G10) activities were derived from optical density and fluorescence measurements obtained after 6 and 8 h of growth, as described previously^18^. The linlog transformation was performed as before^18^, according to:

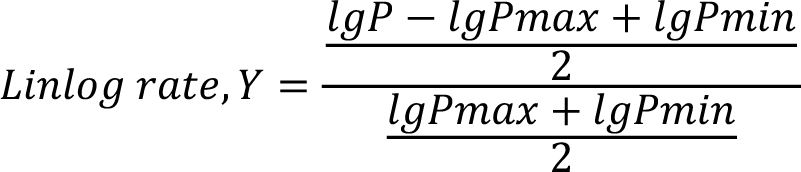

Where;

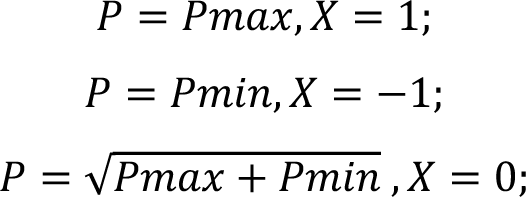

Biosensor expression output and titrations were analysed by fluorescence normalised to cell density (RFU/OD_600)_. Dose response data was fitted with a Hill Function (with variable slope) to extract EC_50_ and Hill slope values using GraphPad (Prism 9). The EC_50_ measurement for library ranking was derived by fitting a Hill function with a standard slope (= 1). JMP was utilised for design of experiments, factor screening and standard least-squares regression analysis. For statistical analysis, all response data was transformed to log10. The corresponding DoE designs were constructed with a definitive screening design (DSD). Factor screening was performed by fitting a two-factor screening model, where the Lenth’s t-ratio from half-normal plots of factor contrast and Lenth’s pseudo standard error (PSE) were used to select significant factors. Those factors with a significant simultaneous p-value (<0.05) were thus used to fit standard least-square regression (SLSR) model(s) of the corresponding response(s). Effect heredity was maintained in these models, and so factors that were significant only in interaction with another factor were also included as individual factors. Response profiler plots were used to analyse and generate factor levels for the desired response characteristics.

### 2.7 Enzyme screening and process optimisation

Esterase and lipase activity towards PET film was assayed using 10 % (w/v) solid loading in 100 mM sodium phosphate buffer pH 7.0 with 25 mg enzyme/g PET. Assays were incubated at 40 °C and 125 rpm for 72 hours before being filtered and stored at -20 °C for TPA determination with the VEC1 biosensor and total product quantification by HPLC. VEC1 biosensor assays were performed as previously described above with the following composition: 450 µl PET hydrolysate, 45 µl ×10 LB, and 5 µl *P. putida* VEC1 preculture. Hydrolysis of PET film by LCC-ICCG at variable temperature and pH was performed using 10 % (w/v) PET solid loading with 10 mg LCC-ICCG/g PET in either 100 mM citrate phosphate buffer (pH 5 or 5.5) or 100 mM Bis-Tris propane (pH 6, 6.5, 7, 7.5, 8, 8.5, or 9). Assays were incubated at either 30 °C, 40 °C, 50 °C, 60 °C or 70 °C and 125 rpm for 72 hours. PET hydrolysate samples were filtered and stored at -20 °C for TPA quantification by the DR1 biosensor and HPLC. DR1 biosensor assays were performed as previously described with 2.5 % (v/v) PET hydrolysate.

### 2.8 Quantification of PET hydrolysis products

400 µl of filtered hydrolysate was appropriately diluted and combined with 400 µl of methanol before injection of 10 µl of sample onto an Agilent 1200 Infinity Series HPLC with a Kinetex 5 µm C18 100 Å column, 250 x 4.6 mM column (Phenomenex^®^) operating at 25 ℃ with an isocratic 70% acetonitrile (0.1% TFA): 30% H_2_O (0.1% TFA) mobile phase at a flow rate of 1 ml/min. Product identification and quantification was carried out using authentic standards.

## 3. Results and discussion

### 3.1 Design of a modular synthetic terephthalic acid biosensor in *P. putida*

To create a fully tuneable terephthalic acid (TPA) biosensors, we aimed to develop a modular genetic architecture with distinct modules for transport, regulation, and output expression. Thus, as an initial step we sought to identify and assemble the required components in order to construct such a system. Several microbial strains including *Comamonas* sp. E6*, R. jostii, P. umsogenesis and I. sakaiensis*, have been reported to possess native TPA metabolism^43,45,46^. In these organisms, the IclR-family transcription factor, TphR, regulates both the TPA catabolic operon *tph* as well as its own transcription from its cognate bidirectional promoter region (P_tph_ promoter)^27,43^. Our previous phylogenetic and genomic context analysis of the TPA transporter, TphC^41^, from *Comamonas* sp. E6 enabled the identification of several novel *tph* catabolic operons across five bacterial families that possed conserved genetic content – including a putative TphR and its respective bidirectional promoter region ^41^ (Fig. 1a, Fig. S1).

**Figure 1.**
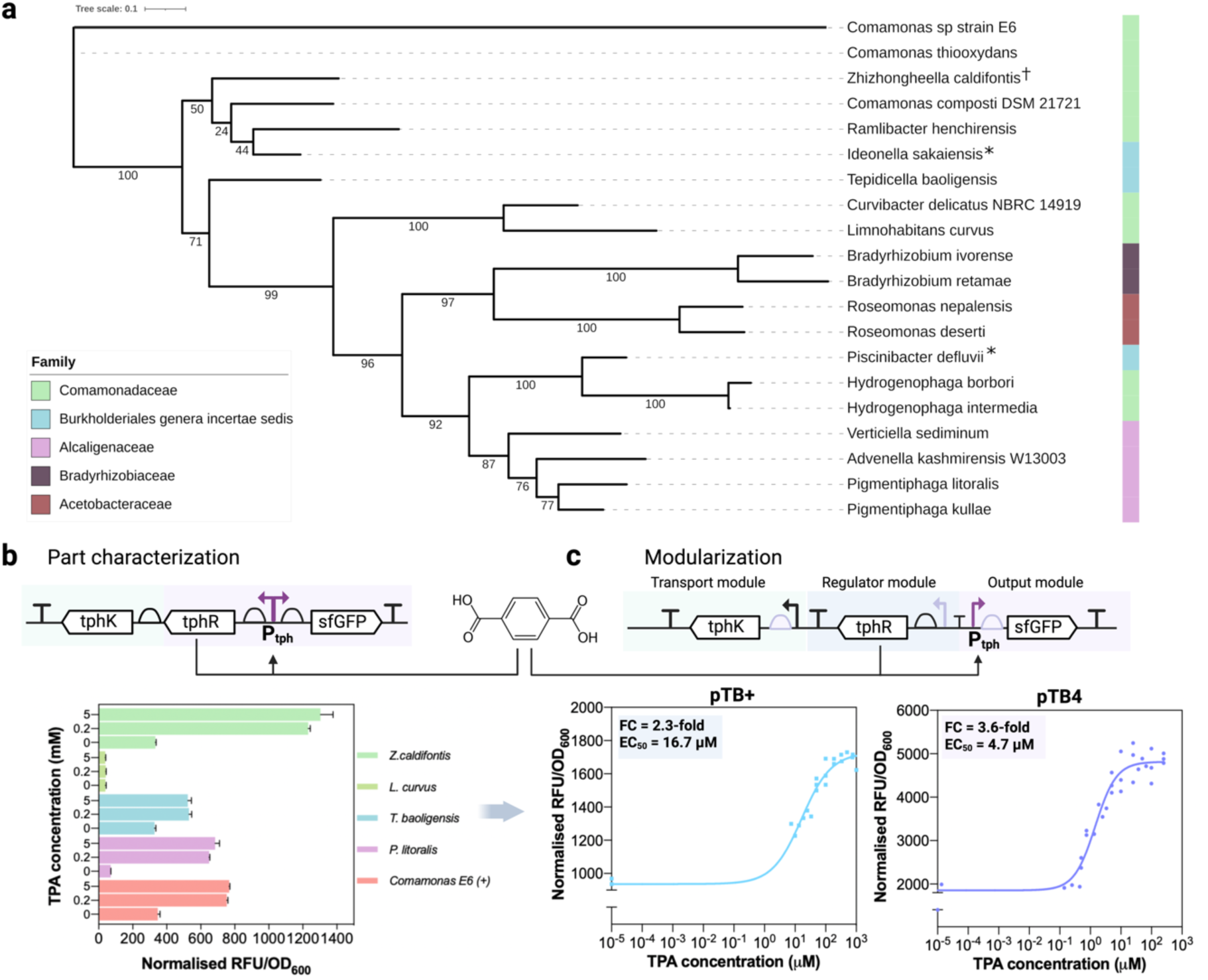
Configuration of a fully modularizable TPA biosensor. (**a)** Phylogenetic tree of the allosteric transcription factor protein TphR. Species names are coloured by taxonomic family. **(b**) Initial biosensor architecture for functional characterization of the putative TphR*-*promoter pairs. The Ptph promoter controls the constitutive expression of *tphR* and *tphK*, and in the presence of TPA, TphR binds to Ptph to initiate transcription of sfGFP. Below are shown the end-point analysis of initial biosensor constructs from a selection of putative homologues. Biosensors were induced with 0, 0.2 and 5 mM of TPA. (**c**) Final modularized TPA biosensor architecture. The PproB and P14A promoters were incorporated to drive expression of *tphR* and *tphK*, respectively.

To investigate the functionality of some of these TphR-promoter pairs as potential biosensors in the host *P. putida*, we selected TphR-promoter pairs from five representative species across a range of habitats and lifestyles, including *Zhizhongheella caldifontis* (a thermophile isolated from a hot spring^47^)*, Limnohabitans curvus* (a psychrotroph isolated from a freshwater lake^48^)*, Pigmentiphaga litoralis* SAS39 (non-type strain of a halotolerant psychrotroph^49^), *Tepidicella baoligensis* (a thermophile isolated from oil well production water^50^), and *Comamonas* sp E6^51^ (isolated from soil). Putative TPA responsive biosensors were constructed wherein the corresponding cognate bidirectional P_tph_ promoter was used to drive expression of both its native *tphR* and the reporter (*sfGFP*) genes. Additionally, since *P. putida* cannot naturally transport TPA, these biosensor cassettes included the gene encoding for the major facilitator superfamily (MFS) TPA transporter from *P. mandelii, tphK* ^45^, downstream of *tphR* within the same operon (Fig. 1b). Each putative biosensor was evaluated for TPA-mediated induction of GFP expression, which revealed two homologue pairs, from *P. litoralis* and *Z. caldifontis* that were functional in *P. putida* and displayed enhanced dynamic range (exhibiting ∼9-fold and ∼4-fold dynamic range, respectively) over sequences from *Comamonas sp. E6* (displaying ∼2.5-fold induction) (Fig. 1b). We verified that these constructs were responsive to TPA and not to other potential PET breakdown products such as BHET or MHET (Fig. S2). Although there was a degree of induction upon addition of MHET, HPLC analysis showed that this was likely due to hydrolysis of MHET to TPA over the course of the induction.

Next, we sought to adapt the architecture of the tested biosensor constructs into a modular format that will be compatible with statistical design approaches. In the tested native configuration, the divergent P_tphC_ and P_tphR_ sequences overlap, which would hinder engineering efforts to tune the expression of the regulation and output modules individually. Thus, to fully decouple expression, we introduced the constitutive P_proB_ promoter^18^ upstream of *tphR* to drive the expression of the regulation module, and insulated it from P_tph_ by the insertion of a terminator and spacer sequence between the two promoters (Fig. 1c). Consequently, the isolated P_tph_ exclusively drives *sfGFP* expression of the output module. Finally, to completely modularise the biosensor architecture, we isolated the transport module by placing *tphK* under the expression of the constitutive P_14A_ promoter^52^ fused to a synthetic G10 RBS sequence^18,53^. The resulting modularised biosensors using sequences from *Z. caldifontis* (hereafter pTB4) and *Comamonas* sp. E6 (hereafter pTB+) yielded fully functional TPA biosensors *and* maintained dynamic ranges from those achieved by the native architectures (Fig. 1c). The pTB4 biosensor displayed increased signal outputs, dynamic range (3.6-fold) and sensitivity (EC_50_ ∼ 5 mM), compared to pTB+, which had lower basal expression but decreased dynamic range (2.3-fold) and sensitivity (EC_50_ ∼ 17 μM) (Fig. 1c). The decoupling was not tolerated by the TphR-promoter pair from *P. litoralis* (hereafter pTB1), as the resulting modularised biosensor construct was not functional.

A terminator was placed between PproB and Ptph to fully decouple Ptph expression. Below are the biosensor dose-response curves of the decoupled pTB+ (left) and pTB4 (right), derived from *Comamonas sp*. E6 and *Z. caldifontis* respectively, induced with a TPA gradient. All results show the ± SD between biological replicates (n = 3).

### 3.2 Refactoring of the biosensor modules and construction of a TphR-responsive promoter library

A crucial first step to exploring the relationship between biosensor design and performance using DoE is the identification of the genetic elements that are expected to influence biosensor response. Further, these factors should be transformed into continuous variables, enabling richer investigation across the defined experimental space of interest.

Optimal biosensor functionality often hinges on achieving a high dynamic range and signal output, which are defined by the concentration and strength of interaction among the regulator and output components. Previously, we optimized these responses in repressor-based systems by adjusting component concentrations at the transcriptional and translational level^18^. However, activator biosensors present a unique challenge due to their interdependent nature; the activity of the output promoter relies directly on factors such as the binding affinity of the regulator to its cognate operator and the concentration of both the ligand transporter and regulator. Consequently, the performance of the promoter cannot be established in isolation from other components. ^5^. As such, we selected four distinct regulation nodes to control the transcription and/or translation of the transporter, transcription factor and output gene: (i) the G10 RBS upstream of the *tphK* transporter (RBS_trans_); (ii) the constitutive P_proB_ promoter regulating *tphR* expression (P_reg_); (iii) the G10 RBS (RBS_out_) upstream of *sfGFP* reporter gene and (iv) the P_tph_ promoter regulating *sfGFP* reporter gene expression (P_out_) (Fig. 2a). To render the RBS_trans_, P_reg_ and RBS_out_ factors into continuous variables, we leveraged previously characterized libraries of the constitutive P_proB_ promoter and the G10 RBS, which were generated and screened in *Escherichia coli* ^18^. Subsequently, we validated the transferability of these libraries to *P. putida* and confirmed their equivalent ranking in *E. coli* (Fig. S3).

**Figure 2.**
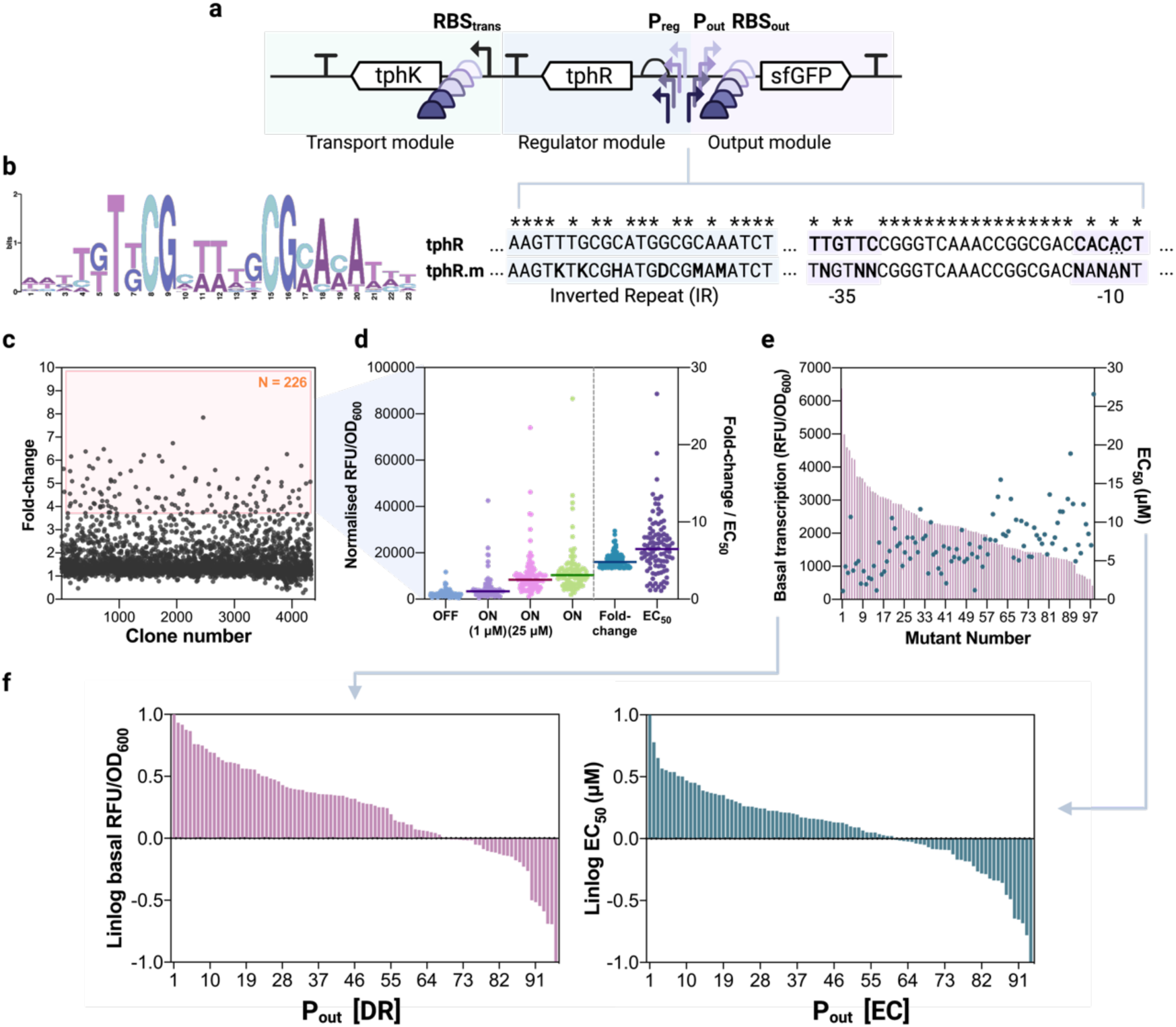
Refactoring of the TPA biosensor and promoter library generation workflow. (**a**) Representation of the biosensor variables selected for refactoring. (**b**) (left) Motif sequence logo of the operator region of Ptph. Generated by MEME without sample correction. (right) Comparison of the wild-type promoter sequence to that of the final mutant design; directed mutations are shown within the operator (blue shaded) and core promoter region (pink shade). (**c**) Fold-change induction of entire promoter library mutants (n » 5,000), and range of biosensor performance (**d**) obtained within the 226 best performing library hits. The shaded line represents the median performance value. The top 226 hits were then re-screened to select a total of (**e**) 100 robust promoter variants that spanned a wide range of basal transcription and sensitivities. (**f**) By differential ranking of these hits, two libraries were obtained based on their strength (left) and sensitivity (right).

However, activator-based biosensors present a unique challenge due to their interdependent nature; the expression of the output module relies directly on factors such as the binding affinity of the regulator to its cognate operator site, interaction between the regulator and RNA polymerase sigma factor, and the concentration of both the ligand transporter and regulator ^5^. Therefore, unlike the other factors, the P_out_ promoter was a discrete sequence, thus for DoE we sought to refactor it into a continuous variable by simultaneously altering the DNA sequences governing activator-operator binding affinity and transcription initiation. A number of studies have previously demonstrated that engineering aTF-operator binding affinity strongly impacts the sensitivity and dynamic range of activator-based system, while changes to the core promoter region further determine the promoter strength and ultimately tune dynamic range ^5,7^. Thus, using a semi-targeted mutagenesis approach for both the operator and core promoter (−35 and -10) regions we can explore a larger portion of the possible design and response space than engineering efforts targeted at either factor individually. To identify which positions of the operator sequence to mutate, we first generalised the operator sequence, building from the previous phylogenetic dataset of *tph* operons to create a general motif consensus sequence of their cognate promoter’s operator region (Fig. 2b; Fig. S1; Fig. S4). This motif overlapped the established *Comamonas* sp. E6 operator motif and was in agreement with a previously described motif for TphR-like IclR regulators (GTNCG-N_5-6_-CGNAC)^44^, with slight differences in peripheral positions (Fig. S4). The motif showed several highly conserved bases as well as some with defined variation between all homologues. From this, we selected 6 positions for mutagenesis, specifically targeting those bases with clearly defined variability for replacement with degenerate nucleotides (Fig. 2b). Within the core promoter region, we targeted the -35 and -10 hex boxes for complete randomisation of three bases in each region (Fig. 2b). These 12 target nucleotide positions correspond a total theoretical promoter library of 489,824 variants. A total sample size of 5,000 (>1%) of these single clonal variants was then assembled in *E. coli* and then screened in *P. putida.* Screening employed an automated workflow where each mutant was assessed for growth and fluorescence in the presence of four distinct ligand concentrations (0, 1, 25 and 1000 mM TPA). Following this initial screen, we selected a total of 226 variants with improved dynamic range relative to the wild-type sequence, which were recharacterized to ensure response robustness (Fig. 2c). Then, from this a total of 100 robust mutants were selected, to reduce library size further. These mutants spanned a wide range of basal and maximal output signals, as well as varied aTF binding affinities (reflected by their varying sensitivities; Fig. 2d). Using this simultaneous semi-targeted promoter engineering approach, we obtained variants that spanned a 17-fold basal transcriptional activity range (from 419 to 6997 RFU/OD_600_) and a 30-fold range of half-maximal effective concentrations, calculated by non-linear regression (EC_50_ ∼1 to ∼30 mM; Fig. 2e).

With the four factors of interest (RBS_trans_, P_reg_, RBS_out_, and P_out_) now having been converted into continuous variables, they were rescaled by a linear-logarithmic (linlog) transformation (eq. 1) into dimensionless variables for DoE. With this rescaling, the minimal and maximal measured outputs for each factor are used to define the extremes of the defined genetic space, and correspond to rescaled values of -1 and +1, respectively, while the geometric average from both corresponds to the central level 0. The linlog rescaling of the P_reg_, RBS_trans_ and RBS_out_ factor libraries was done according to protein synthesis rate (Fig S2), which correlated well with the ranking previously observed in *E. coli* ^18^. While the P_out_ factor library was ranked separately according to both promoter strength (DR library) or sensitivity (EC library). This ranking of the P_out_ library based on two different criteria was performed to learn whether enriched knowledge and, eventually, distinct biosensor performances could be attained by differentially ranking the same genetic context (Fig. 2f). As such, the maximal and minimal P_out_ basal transcriptional strengths were set as +1 and -1 for the promoter strength ranked (DR) library, while the maximal sensitivity (lowest EC_50_) was set to -1, and minimal (highest EC_50_) set to +1 for the sensitivity ranked (EC) library.

### 3.3 Differential promoter ranking leads to distinct design space exploration

Considering the number of selected factors and the size of their corresponding libraries, a total of more than 1×10^6^ combinations (22 x 22 x 22 x 100) would be necessary to fully sample the entire biosensor design space. Instead, we employed a Definitive Screening Design (DSD) that would allow us to fractionally sample this design space with the aim of identifying the main factor effects and interactions while minimizing experimental effort. DSD employs a three-level screening approach and permits estimation of curvature effects, ideal for direct screening and optimisation efforts, as opposed to other factorial two-level designs commonly used in DoE, such as Plackett−Burman designs^24,26^. Here, using DSD we were able to sample the design space to just 17 discrete combinations, including four additional designs to ensure robustness of the factor significance. Two DSD experiments were performed for the separately ranked P_out_ promoter library (hereafter named the *DR* and *EC* models). Constructs were assembled according to the 17 genetic designs for each model, as shown in Table S1, where each factor was set to the specified design level (Fig. 3a). We note, however, that after continuous efforts, one of the experimental designs – wherein every component was set to its maximal expression (run #13, Table S1) – could not be assembled despite multiple attempts. This can likely be attributed to the toxicity and high level of burden resulting from the expectedly strong expression of every biosensor module, especially the transport module encoding the membrane transport protein, TphK. After transformation of *P. putida* with the assembled plasmids, we assessed their performance by performing TPA titrations across an increasing concentration gradient, ranging from 0 to 0.5 mM. Dose-response measurements were then fit with a Hill function that permitted careful analysis of signal outputs (ON and OFF), dynamic range (ON/OFF), sensitivity (EC_50_) and curve steepness (Hill slope), which constituted the model responses.

**Figure 3.**
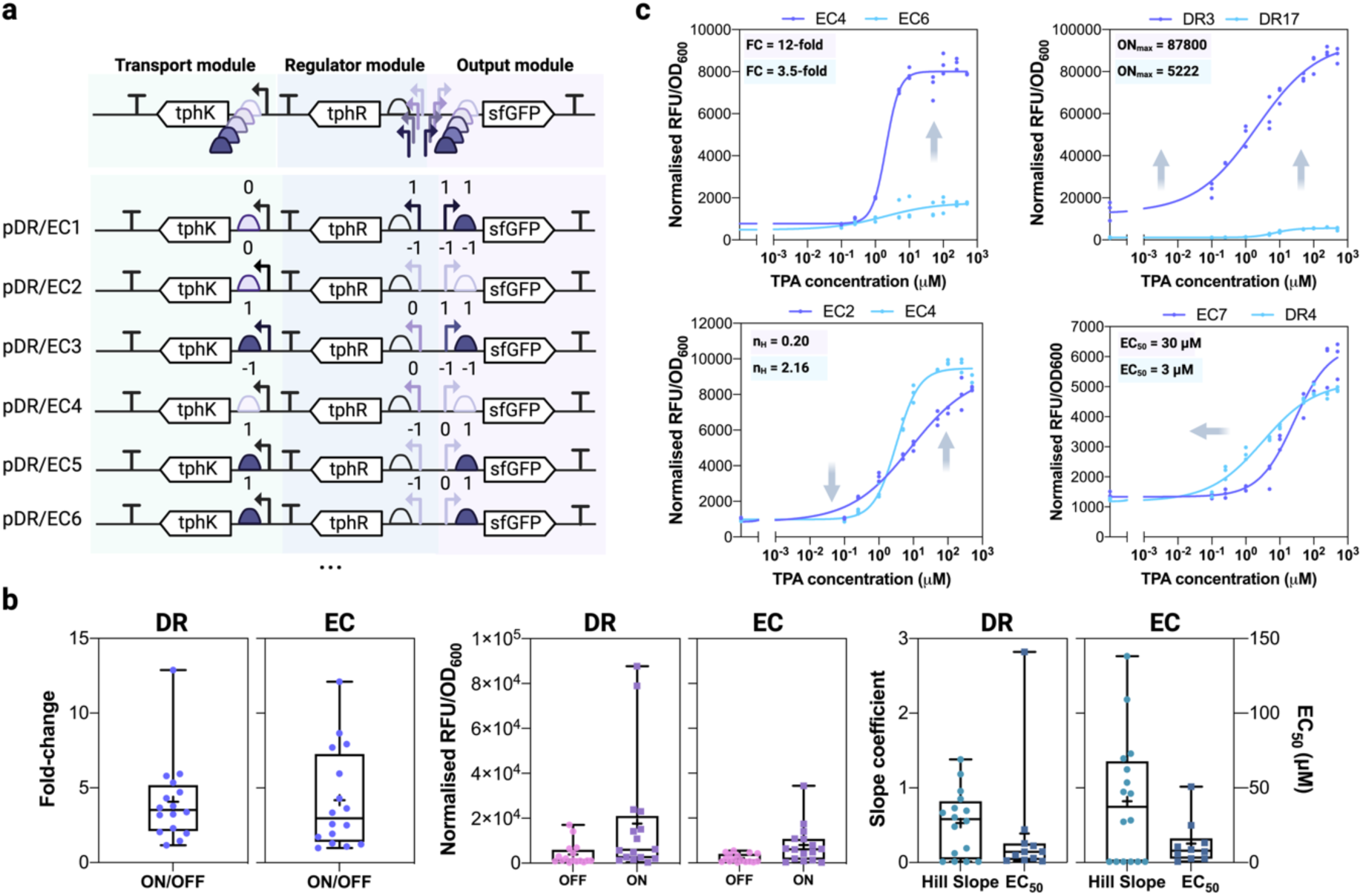
Experimental DoE trials. (**a**) Genetic configuration of the experimental designs conforming to the specified definitive screening design. Six exemplar designs are shown, all designs can be found in Table S1. (**b**) Range of biosensor performances (dynamic range, signal outputs and slope and sensitivity parameters) obtained from both experiments. For each individual graph, the left panel corresponds to DR ranking and right panel to EC ranking. Middle box line represents the median value obtained, while the + sign corresponds to the mean value. (**c**) Examples of different constructs that exhibit altered response of a single performance (those constructs exhibiting similar remaining performances were chosen to best illustrate the difference of the given response). All results show the ± SD between biological replicates (n = 3).

The resulting constructs exhibited a broad range of performances between the two DSD experiments (Fig. S5, Fig. 3b; Table S2, Table S3). Both experiments contained constructs that exhibited increased maximal signal output – over 18-fold (from 4892 to 87800 RFU/OD_600_) – and dynamic range – 4-fold (3.6-fold to 13-fold) – compared to the original pTB4 construct. Moreover, the tested biosensors ranged in sensitivity across three orders of magnitude (from EC_50_ in the nM to mM range) and produced hill slopes exhibiting both digital and analogue responses (Fig. 3c). Interestingly, the best performers from both models possessed similar performance in terms of dynamic range, while they varied in their slope, sensitivity, and signal outputs (Fig. 3c). In contrast, the poorest performers were barely responsive, highlighting the significant advantage of using a combinatorial based approach for biosensor design. As expected, we observed enrichment of different response performances from the DR and EC models. As the coverage of promoter transcriptional strength was greater in the DR model, constructs with high transcriptional output both at the OFF and ON responses were enriched here (maximal DR ON = 87,800 RFU/OD_600_ vs. maximal EC ON = 37,880 RFU/OD_600_) (Fig. 3b). In contrast, the EC model demonstrated increased variability of hill coefficients (steepness) and higher sensitivity (Fig. 3c). These differences denote that simultaneous analysis of both P_out_ design spaces could thus provide a greater exploration of the causative effects and underlying interactions that lead to each response, and which might ultimately lead to achieving a desired dose-response.

To understand the main effects governing biosensor performance in both DSD experiments, the chosen factors affecting the five distinct responses (ON, OFF, ON/OFF, Hillslope and EC_50_) were statistically evaluated by analysis of variance (ANOVA) and two-level screening analysis. Based on mean value distributions and half-normal probability plots, a simultaneous p-value can be assigned to investigate factor significance on each response (Fig. S6-S7). Therefore, changing the level of those factors with the largest primary effects is expected to produce the greatest variance. The results obtained indicated differences in factor significance upon the overall dose-response between both experiments (Fig. S5; Fig. S6-S7). Full detailed results on factor significance for each response and experiment can be found in Fig. S6 and Fig. S7. As both rankings aimed to explore the impact of the different effect extremes from P_out_ (strength and sensitivity), differing factor significance validates the successful exploration of the effects emerging from such extremes, which would be impossible to assign simultaneously.

### 3.4 Statistical modelling enables the exploration of combined biosensor response conditions

To investigate the resulting significant effects on biosensor performance, we next built standard least-squares regression (SLSR) models using the factors found to be significant (p < 0.05) for each response (Fig. S6-S7). The resulting SLSR models accurately described the relationships within the data for both experiments and all responses (p < 0.0001; min R² = 0.94; Fig. S8-S9). Exploring DSD designs with SLSR, we aimed to illustrate how changes to individual biosensor components affect the dose-response curve in both DR and EC rankings. This approach can help obtain a prediction of the resulting phenotypic landscape of the five responses simultaneously. Altering a specific factor (e.g., changing P_reg_ promoter strength) can have effects on multiple responses, and thus defining these response trade-offs is essential to understand what changes lead to the required performance.

Both the DR and EC models provide valuable insights into optimising the ON/OFF response of the constructed biosensors. In the DR model, an almost linear relationship between P_out_ and RBS_out_ and the ON/OFF response is observed (Fig. 4a). Simultaneously increasing both of their strength results in greater output (ON), but with a less pronounced effect on OFF. This leads to greater fold-difference between ON and OFF at increased levels of P_out_ and RBS_out_. Overall, changing P_reg_ has a null impact on ON/OFF as both ON and OFF levels are equally impacted. Specifically, at high P_reg_ both ON and OFF levels drop, suggesting that excessive TphR can interfere with stable aTF-promoter activation or cause tetramer aggregation^54^. In contrast, tuning P_reg_ from low to mid-range does not have a strong impact upon ON or OFF, indicating that lower TphR levels satisfy the transcription initiation requirements of P_out_ (Fig. 4a). Together, this leads to an optimal predicted configuration from the DR ranking to afford a biosensor with enhanced dynamic range: a P_out_ promoter with a high transcriptional strength should be paired with a strong RBS_out_. The level of regulator should then be fine-tuned to ensure optimal interaction with P_out_. This configuration can be exemplified by DR1, the closest design in accordance with the optimal predicted profile (RBS_trans_ (0); P_reg_, P_out_ and RBS_out_ (+1); Table S2, Fig. 4c).

**Figure 4.**
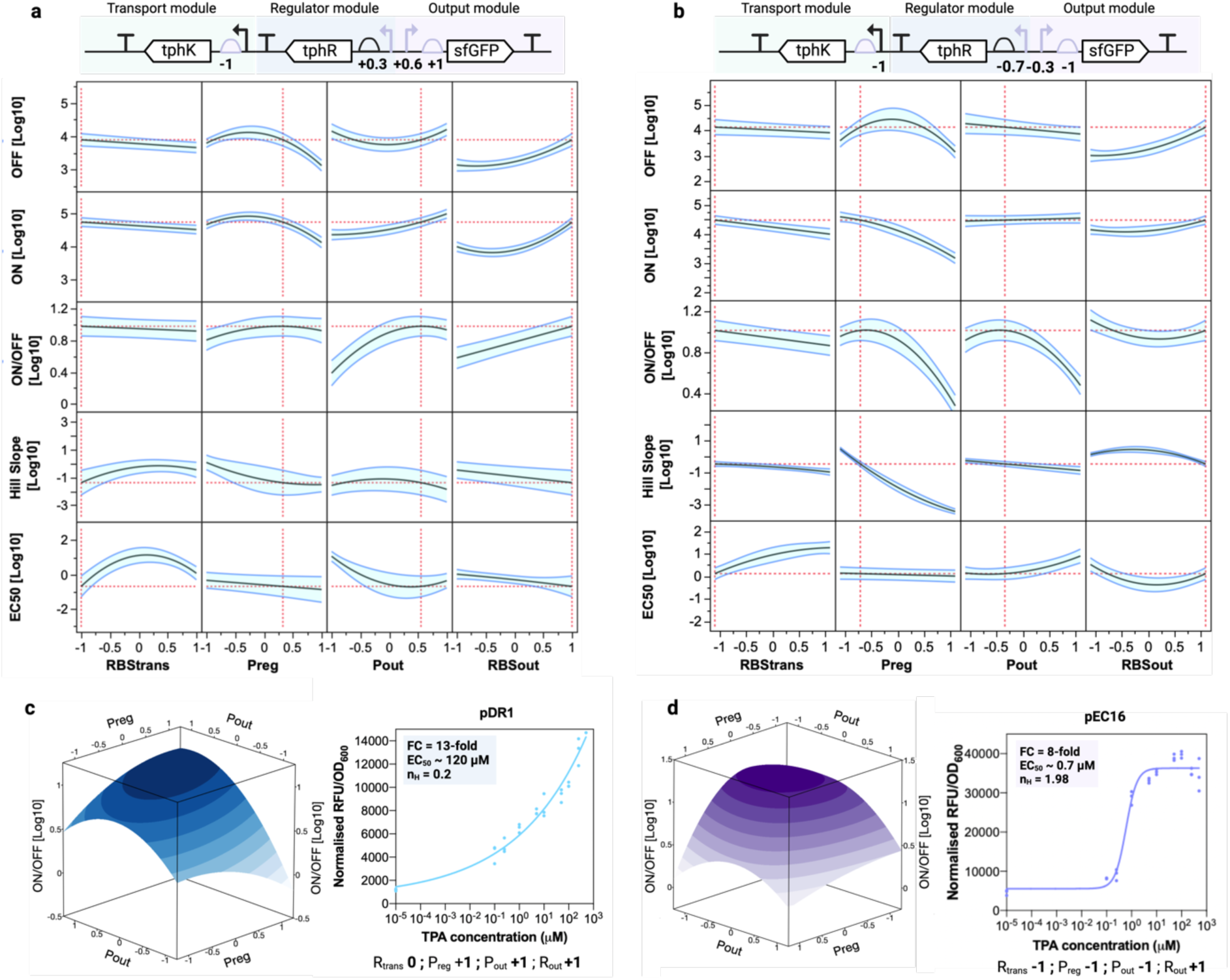
Modelling biosensor performance. (**a**) DR ranked library - prediction profiler of SLSR model displaying optimal design for improved dynamic range and signal output. (**b**) EC ranked library - prediction profiler of SLSR model displaying optimal design for improved dynamic range and signal output. (**c**) DR - Surface plot of dynamic range between Preg and Pout also shows non-linear effect and a global optimum (left) and closest experimentally validated biosensor construct, pDR1, to the optimal indicated by SLSR model. (**d**) EC - Surface plot of dynamic range between Preg and Pout also shows non-linear effect and a global optimum (left) and closest experimentally validated biosensor construct, pEC16, to the optimal indicated by SLSR model.

The EC ranking, conversely, provides alternative insights into promoter-driven dynamic range performance variability. The effect from P_out_ in ON/OFF seen in this ranking is reversed, meaning that when ranking the promoter by EC_50_, an increase in P_out_ (weaker affinity) leads to a drop in OFF and therefore ON/OFF; this slight inverse effect can also be observed from the ranking data directly (Fig 2e). Additionally, increased output signal (ON) in this ranking is thus predominantly driven by the output translational strength (RBS_out_)(Fig. 4b). Increasing RBS_out_ has a greater impact upon OFF, meaning that between 0 and +1, ON is limited by another component in the system, in this case seen by P_reg_ (Fig. 4b). A slight trade-off between maximal ON and ON/OFF is therefore observed by the EC model: in order to match P_out_ and display the maximal ON, higher ON/OFF is achieved only if both P_reg_ and P_out_ are kept at low levels. However, slightly higher ON/OFF can be achieved if RBS_out_ is kept low, but with a significant drop in maximal ON achieved (Fig. 4b). These insights can be exemplified by the closest optimal example EC16, which achieved both good dynamic range and maximal signal output (RBS_trans_ (−1); P_reg_, P_out_ (−1) and RBS_out_ (+1), Fig. 4d).

Optimising dynamic range and signal-to-noise ratio is generally a performance pre-requisite for transcriptional biosensors. However, for other biosensor responses, including sensitivity and hillslope, maximising these responses might also be the goal. Within the context of this DoE framework, the user may need to fine-tune these responses and adjust the objective functions according to their specific requirements (e.g., maximizing or minimizing the Hill slope). Therefore, here we chose to explore how both sensitivity and steepness vary at the predicted optimal ON/OFF settings for both DR and EC rankings. For both models, the maximal achievable sensitivity (low EC_50_) is observed at the proposed settings shown for maximal ON/OFF (Fig. 4a,b). Evidently, as outlined above, sensitivity was inversely impacted by P_out,_ when ranked by either DR or EC. In the EC model, a slight further increase in sensitivity is shown to be achievable by lowering RBS_out_, but at the expense of lowering ON and OFF (Fig. 4b). Furthermore, both models display complementary trade-offs between sensitivity and the achieved curve steepness. To achieve changes in curve slope, previous efforts have necessitated targeted protein engineering approaches and other additional multipart regulatory mechanisms, such as aTF decoy binding “sponges”^55^ or bistability feedback networks, and normally result in parallel effects on other biosensor responses^2^. In the DR model, within the optimal ON/OFF levels defined above, the curve steepness is seen to lay within a linear range (Fig. 4a). Tuning the regulator and the available ligand concentration can achieve some minor changes in curve steepness while maintaining high dynamic range, but at the expense of a decrease in sensitivity (Fig. 4a). Conversely, for the EC ranking, the optimal settings predict a steeper hillslope, and more significant effects are observed for this response, whereby increasing P_reg_, and to a lesser extent P_out_ and RBS_out_, results in reduction on hill slope (less digital)(Fig. 4b).

### 3.5 Proposed model validation yields custom high-performing TPA biosensor

Using DSD for both DR and EC rankings provided complementary insights into biosensor performance optimization. We were able to identify distinct non-linear effects and causative factors between the five explored responses, which can direct biosensor engineering efforts without the need for additional experimental data. The results validated that the design space obtained from tuning the P_out_ promoter for sensitivity (EC model) can be used to inform the development of biosensors with high dynamic range, sensitivity, and steep digital dose-response for both high and lower signal outputs. Such a biosensor could find application in yes-or-no screening assays. On the other hand, the design space obtained from tuning the P_out_ promoter for strength (DR model) can inform the development of biosensors with optimal signal-to-noise ratios and analogue dose responses. Such a biosensor system could find application in the accurate distinction between close analyte concentrations over a wide output range.

To test the rigour of the framework developed here for custom TPA biosensor performance engineering, we constructed the optimal trial proposed by the EC model wherein the biosensor responses were treated as a combined object function set for obtaining maximal signal outputs (ON), sensitivity (EC_50_), steepness (nH), and high dynamic range (ON/OFF) simultaneously. The factor settings for said biosensor were as follows: RBS_out_ was set to the highest level (+1), P_reg_ was set to -0.7 and the level of P_out_ was set to the advised mid-lower point (−0.3), while RBS_trans_ was maintained at -1 (Fig. 5a). The resulting construct (hereafter VEC1) gave the best combined performance of all tested biosensors so far, with a sensitivity of EC_50_ = 0.7 uM, maximal ON = 37156 RFU/OD_600_, dynamic range of 13.2-fold, and curve steepness of nH = 2.7 (Fig. 5). Compared to the original pTB4 biosensor, VEC1 demonstrated an 8-fold improvement in output signal, 7-fold increase in sensitivity, 3.7-fold higher dynamic range, and a highly digital hill coefficient (the highest nH seen from all constructs (from nH = 1.98 in pEC16 to nH = 2.5 in pVEC1)(Fig. 5). This construct successfully validates the proposed EC model, with confidence that the optimal combined configuration of this biosensor has been achieved. Moreover, this suggests that experimental space and potential knock-off effects had been efficiency explored, which is particularly crucial when optimising more than one response simultaneously. For instance, a highly active TPA biosensor was previously developed in the bacterium *C. thiooxidans* strain S23, which can natively transport and metabolise TPA^27^. The initial biosensor version exhibited an excellent ON/OFF response up to 25-fold but with lower sensitivity (in the micromolar range). Subsequent metabolic engineering efforts to improve its sensitivity resulted in biosensor capable of responding to TPA in the nanomolar range. However, this enhancement in sensitivity came with a knock-on effect on the dynamic range, reducing the ON/OFF response to below 5-fold^27^. The biosensor framework presented here demonstrates the potential of using DoE to maximize all responses while avoiding these knock-on effects and extensive metabolic engineering efforts, thereby yielding a TPA biosensor with the best combined activities achieved to date ^27,29,56^.

**Figure 5.**
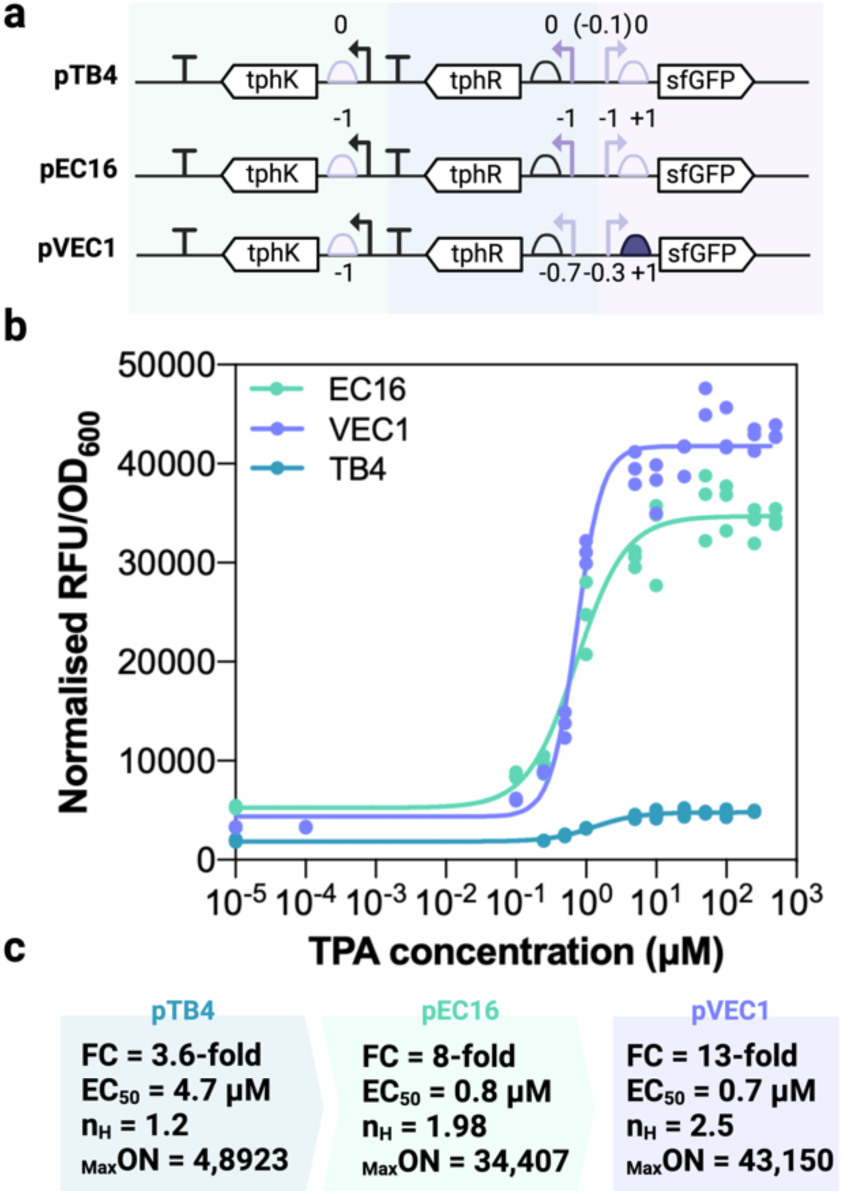
Optimisation and validation of EC-model based biosensor VEC1. (**a**) Shown are the diagrams of the biosensor configurations for the original (TB4), intermediate (EC16) and final optimal (VEC1) biosensor construct. In the validation construct VEC1, the level of Pout and Preg were set to the predicted optimal configuration as dictated by the EC model. (**b**) The resulting dose-response curves of the biosensors show optimisation to the signal-to-noise ratio, dynamic range, sensitivity and hillslope of the biosensor, as illustrated by their (**c**) performance metrics. All results show the individual points of three biological replicates (n = 3).

### 3.7 Application of the TPA biosensors in primary and secondary PETase activity screening

The convenience and advantage of biosensors as a metabolite detection system has resulted in a significant growth in their application as a tool for optimizing the metabolic flux of microbial bioproduction strains or for high-throughput screening of strain/enzyme libraries for desired phenotypes^12,57^. Often, these roles require specific ligand response profiles (sensitivity, operating range, dynamic range, specificity, etc) for optimal application performance (Fig. 6a). Enzyme mediated PET plastic recycling is believed to be superior to mechanical recycling as it requires less energy input and retains the material value of the polymer^58,59^. Towards this, discovery and development of more efficient PET depolymerising PETase, cutinase and hydrolase enzymes is of great biotechnological interest. Evaluating the activity of these enzymes often relies on low-throughput chromatography-mediated quantification of hydrolysis products. Recently, however, an enzyme coupled luciferase reporter system in *E. coli* and a the optimal TphR-P_tphC_ biosensor expressed in *Comamonas thiooxidans*, were applied to the detection of TPA released from enzyme hydrolysed PET ^28^, demonstrating the potential of TPA responsive biosensors for the semi-quantitative high throughput screening PET active enzymes. Thus, we explored the application of the developed TPA-responsive biosensors for two distinct enzyme activity screening applications (Fig. 6a).

**Figure 6.**
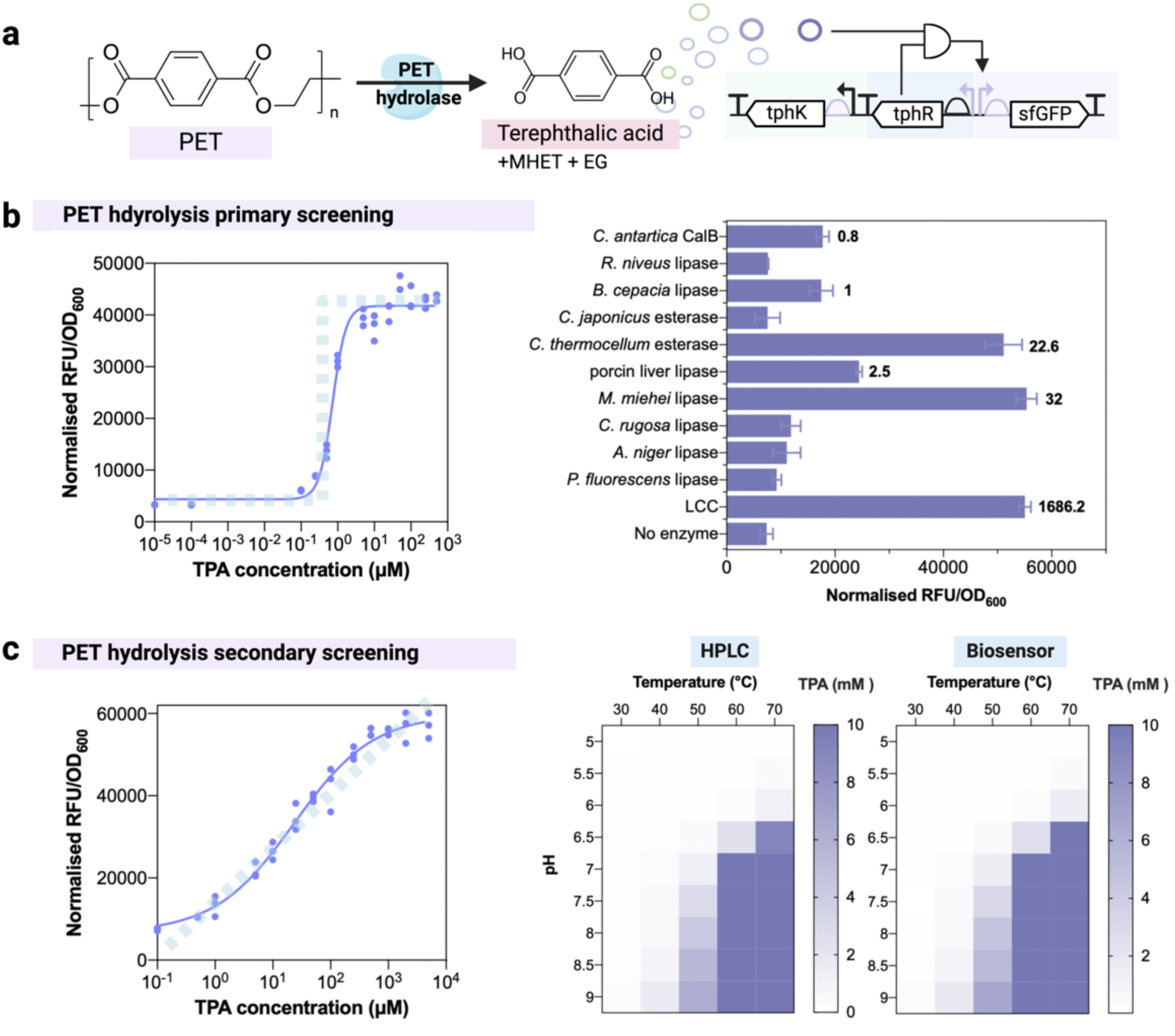
Deployment of the developed biosensors in primary and secondary PET hydrolase screening applications. (**a**) After enzymatic hydrolysis, PET plastic is depolymerised to TPA, MHET and EG. TPA biosensing can therefore find applications in the discovery and development of more efficient PET depolymerising PETase, cutinase and hydrolase enzymes. (**b**) The application of the VEC1 biosensor is optimal for primary YES/NO enzymatic screening, as this biosensor displays a highly digital, sensitive and strong TPA induced response (left). VEC1 was thus used to screen a selection of esterases and lipases for PET hydrolysis activity (right). The bold numbers next to each column display the corresponding TPA concentration measured by HPLC (Fig. S10). (**c**) The application of the DR1 biosensor is optimal for secondary enzyme screening, as this biosensor is responsive over an extended range displaying a highly analogue standard curve (left). DR1 was thus used to screen enzyme process conditions of the highly active LCC-ICCG cutinase across a temperature and pH gradient (right), which showed excellent accordance with HPLC quantification. All results show the individual points of three biological replicates (n = 3)

Firstly, we demonstrated the application of the VEC1 biosensor for primary screening, the objective of which is to provide a rapid and binary response for any basal activity of the desired function. As such the screening method is required to be highly digital, sensitive to the substrate concentration, and greatly responsive by displaying high signal-to-noise ratio - demonstrating clear binning of active and inactive candidate enzymes. Such screening is often employed for enzyme discovery. Here we screened a selection of esterases and lipases from *C. thermocellum*, *C. japonicus*, *P. fluorescens*, *A. niger*, *C. rigosa*, *M. miehei*, *B. cepacia*, *R. niveus*, and *C. antarctica* for their ability to hydrolyse the PET ester linkage and release TPA. The engineered cutinase, LCC-ICCG, was used as a positive control for PET depolymerisation^60^. Amorphous PET film was incubated with purified enzyme at 40 °C and pH 7 for 72 hours before the supernatant was harvested and incubated with the *P. putida* VEC1 reporter strain for fluorescence mediated detection of TPA. It was found that the VEC1 reporter strain was capable of distinguishing between those enzymes which released TPA from PET and those that did not with high fidelity (Fig. 6b Fig. S10). Further, hydrolysate containing TPA concentrations as low as 800 nM elicited a fluorescent response above the baseline, demonstrating the sensitivity of VEC1 biosensor is maintained in an application setting.

Secondly, we demonstrated the application of the DR1 biosensor to secondary screening, the aim of which is to distinguish between closely related activity profiles. As such, the screening method is required to be responsive over an expanded range of analyte concentrations and display a linear dose response slope that can be used for exact quantification (Fig. 6c). Such screening is often employed to distinguish between engineered enzyme variants or homologues with closely related activity profiles, or to optimise the environmental and process conditions of an enzymatic reaction^12,57,61^. Here we opted to screen the PET depolymerisation activity of the LCC-ICCG cutinase across a temperature and pH gradient of 30-70 °C and 5-9, respectively. PET film hydrolysis assays were harvested after 72 hours and incubated with the *P. putida* DR1 reporter strain for fluorescence mediated quantification of released TPA. The DR1 biosensor was selected as it displayed a broad operational range (of >3 orders of magnitude) and a highly analogue response (n_H_ = 0.2), and in this experimental setup afforded a fold-change of 8-fold over the TPA range tested (Fig. 6c). Biosensor mediated quantification of TPA present in PET hydrolysates was performed using the equation for the line of the linear range of the dose response curve. Here it was found that the DR1 calibration curve was able to accurately determine the concentration of released TPA as compared to HPLC, with fluorescence quantification typically being within 15% of HPLC determined values (Fig. 6c; Fig. S11). Collectively these exemplar applications effectively demonstrate that biosensors can provide a rapid, low cost, high-throughput, and built-for-purpose alternative to the common semi-quantitative and quantitative PET degradation screening strategies.

## 4. Conclusions

In summary, this study proposes using DoE as a framework for achieving tailored activator-based biosensor performance. The dual promoter refactoring approach allowed simultaneous and enriched investigation into the causative effects from the core and operator region library upon each specific response effect. Furthermore, by employing a fractional sampling approach via DSD, these learnings were possible to attain with a significantly reduced experimental design space. Specifically, employing only 32 unique genetic designs between two distinct experiments, which represent merely 0.003% of the total design space investigated, we were able to identify key genetic effects that ultimately led to functional TPA biosensors with increased maximal signal outputs over 18-fold (from 4892 to 87800 RFU/OD_600_) and dynamic range by about 4-fold (3.6-fold to 13-fold). Moreover, the obtained biosensors spanned sensitivities covering three orders of magnitude and hill slopes ranging a large cooperative binding space (from nH <1 up to >2). The experimental space covered in this single experiment was enough to suggest a tailored biosensor configuration to simultaneously maximise all five responses. This was successfully validated with the construction of a single VEC1 construct, which gave the best combined performance of all biosensors tested, validating the design framework as a time and cost-efficient methodology to guide engineering for a specific response combination. The same methodology employed for VEC1 could be followed for any response combination needed, and which can lead to the development of panel of tailored biosensors for diverse applications.

Following the design rules elucidated here will aid the development of biosensors with custom performances necessary for distinct applications in plastic PET valorisation, bioremediation, assimilation and/or bioproduction^35,37–39,57^. Using the same starting point and genetic library elements, this framework has successfully led to the development of two novel analytical methodologies for PET hydrolysis screening. The identification of PET-degrading enzymes generally relies on enrichment cultures and homology-based and phylogenetic methods ^62–66^. These strategies have led to the discovery of numerous enzymes capable of degrading PET, polyurethane (PUR), and other polyamide (PA) oligomers, and has significantly expanded our understanding of the catalytic mechanisms of plastic hydrolysis ^59,66^. However, there remain substantial gaps in our knowledge regarding the biological functions, diversity, and evolutionary pathways of these enzymes and/or pathways. Additionally, there is a need to enhance the diversity of known plastic-active enzymes by identifying genuinely novel enzymes, with distinct folds, through the detection of their reaction products. This is a vital area of research that promises to deepen our understanding and broaden the spectrum of plastic-degrading enzymes. By applying statistical modelling here, we were able to obtain biosensor with tailored responses without needing extensive engineering of the activator protein, and without previous knowledge on the underlying activation mechanism. The framework proposed here is therefore ideal to follow when building new uncharacterized systems for novel inducers and/or microorganisms, facilitating robust data-driven decision making.

## Acknowledgements

We would like to thank The Henry Royce Institute for Advanced Materials (funded through EPSRC grants nos. EP/R00661X/1, EP/S019367/1, EP/P025021/1 and EP/P025498/1) for access to their facilities. We would also like to thank Alejandro Marquiegui for his valuable help with sorting the library data and to Mark Dunstan for assisting in automation efforts.

## Author contributions

G.A and N.D conceived the project. G.A, M.C and T.B. designed and carried out the experiments. G.A wrote the manuscript. All the authors read and approved the final manuscript.

