## Supplementary Information for "Tuning the performance of a TphR-based terephthalate biosensor with a design of experiments approach"

**Contents**

**Table S1.** Definitive Screening Design proposed by JMP platform.

**Table S2.** Results obtained from dose-response curves of all respective EC ranking constructs.

**Table S3.** Results obtained from dose-response curves of all respective EC ranking constructs.

**Table S4.** Plasmids used in this study.

**Table S5.** List of primers and oligos used in this study.

**Figure S1.** Distribution of TSSs and hex-boxes inferred using the extended thermodynamic model within *tph* and *tph*-like promoters.

**Figure S2.** TPA Biosensors treated with PET breakdown products.

**Figure S3.** Constitutive ProB promoter and RBS G10 libraries activity in P. putida.

**Figure S4.** Distribution of putative TphR binding sites within *tph* and *tph*-like promoters.

**Figure S5.** Full experimental trials data for DR and EC experiments.

**Figure S6.** Factor screening and selection from DR experiment.

**Figure S7.** Factor screening and selection from EC experiment.

**Figure S8.** Standard Least Squares Regression model performance for the DR experiment.

**Figure S9.** Standard Least Squares Regression model performance for the EC experiment.

**Figure S10.** Esterase and lipase activity towards PET film.

**Figure S11.** Biosensor- versus HPLC-mediated quantification of TPA.

**Table S1. Definitive Screening Design proposed by JMP platform.** The respective DSD vectors were constructed according to this design for both the DR and EC rankings.

| DR/EC Run | $RBS_{trans}$ | $P_{reg}$ | $P_{out}$ | $RBS_{out}$ |
|---|---|---|---|---|
| **1** | 0 | 1 | 1 | 1 |
| **2** | 0 | -1 | -1 | -1 |
| **3** | 1 | 0 | 1 | 1 |
| **4** | -1 | 0 | -1 | -1 |
| **5** | 1 | -1 | 0 | 1 |
| **6** | -1 | 1 | 0 | -1 |
| **7** | 1 | -1 | -1 | 0 |
| **8** | -1 | 1 | 1 | 0 |
| **9** | 1 | 1 | -1 | -1 |
| **10** | -1 | -1 | 1 | 1 |
| **11** | 1 | -1 | 1 | -1 |
| **12** | -1 | 1 | -1 | 1 |
| **13** | 1 | 1 | 1 | 1 |
| **14** | -1 | -1 | 1 | -1 |
| **15** | 1 | 1 | 1 | -1 |
| **16** | -1 | -1 | -1 | 1 |
| **17** | 0 | 0 | 0 | 0 |

**Table S2. Results obtained from dose-response curves of all respective EC ranking constructs.**

| | OFF | OFF ± SD | ON | ON ± SD | ON/OFF | ON/OFF ± SD | $EC_{50}$ (mM) | Hill Slope |
|---|---|---|---|---|---|---|---|---|
| **DR1** | 1179.3 | 80.757 | 15138 | 379 | 12.881 | 1.0264 | 153.15 | 0.1587 |
| **DR2** | 804.18 | 116.31 | 2583.4 | 206.62 | 3.2347 | 0.23 | 1.842 | 0.5585 |
| **DR3** | 14086 | 4441.5 | 87740 | 3631.5 | 6.7892 | 2.7163 | 2.13 | 0.4833 |
| **DR4** | 1143.3 | 108.95 | 4899.3 | 87.865 | 4.3068 | 0.3404 | 21.07 | 0.4106 |
| **DR5** | 2047 | 175.39 | 10899 | 388.87 | 5.3428 | 0.3391 | 2.432 | 0.5336 |
| **DR6** | 454.5 | 36.291 | 1663.7 | 38.488 | 3.6713 | 0.2055 | 3.136 | 1.5665 |
| **DR7** | 1187.6 | 122.89 | 2708.4 | 127.31 | 2.2944 | 0.2199 | 114 | 0.9403 |
| **DR8** | 1009.9 | 71.609 | 2055.4 | 83.313 | 2.0381 | 0.0632 | 0.1 | 0.01 |
| **DR9** | 312.13 | 34.911 | 358.66 | 5.4402 | 1.1578 | 0.1183 | 2.7 | 0.01 |
| **DR10** | 17007 | 2529.9 | 78998 | 5825.2 | 4.7048 | 0.694 | 1.031 | 0.5879 |
| **DR11** | 6422.2 | 953.08 | 23879 | 787.35 | 3.7846 | 0.6781 | 15.11 | 0.5136 |
| **DR12** | 2920.8 | 132.41 | 5703.4 | 233.52 | 1.9552 | 0.1151 | 7.5 | 0.01 |
| **DR14** | 6851.6 | 805.09 | 23021 | 1406.1 | 3.3775 | 0.2381 | 0.4937 | 1.207 |
| **DR15** | 906.74 | 65.629 | 1305.9 | 57.379 | 1.4424 | 0.048 | 0.2 | 0.01 |
| **DR16** | 4486 | 175.04 | 14244 | 1254.1 | 3.1771 | 0.3065 | 10.13 | 1.579 |
| **DR17** | 1089.4 | 36.291 | 5222.4 | 721.83 | 4.915 | 1.3236 | 7.204 | 0.9196 |

**Table S3. Results obtained from dose-response curves of all respective EC ranking constructs.**

| | OFF | OFF ± SD | ON | ON ± SD | ON/OFF | ON/OFF ± SD | $EC_{50}$ (mM) | Hill Slope |
|---|---|---|---|---|---|---|---|---|
| **EC1** | 266.89 | 1.7171 | 289.56 | 17.743 | 1.0851 | 0.0702 | 109.14 | 0.01 |
| **EC2** | 733.65 | 15.332 | 6346.6 | 145.75 | 8.6541 | 0.3056 | 8.543 | 0.4957 |
| **EC3** | 5498.2 | 65.9 | 10650 | 169.1 | 1.9372 | 0.046 | 1.048 | 0.01 |
| **EC4** | 652.92 | 35.418 | 7886.5 | 23.938 | 12.103 | 0.6819 | 3.225 | 2.167 |
| **EC5** | 1773.9 | 82.551 | 14072 | 1335.7 | 7.9368 | 0.7358 | 5.324 | 0.5336 |
| **EC6** | 503.5 | 61.392 | 1751.2 | 37.547 | 3.5054 | 0.3381 | 3.136 | 1.5665 |
| **EC7** | 1388.7 | 111.6 | 5941.4 | 616.71 | 4.297 | 0.5848 | 23.42 | 0.8355 |
| **EC8** | 98.589 | 3.654 | 173.14 | 116.06 | 1.7465 | 1.1532 | 4.92 | 0.01 |
| **EC9** | 4205 | 208.86 | 4098.3 | 327.68 | 0.9775 | 0.1085 | 104.9 | 0.01 |
| **EC10** | 4486 | 91.005 | 14244 | 1254.1 | 3.1771 | 0.3065 | 16.27 | 0.9144 |
| **EC11** | 738.63 | 12.021 | 1910.9 | 91.026 | 2.5873 | 0.1249 | 6.362 | 1.099 |
| **EC12** | 3705.2 | 228.28 | 4789.5 | 205.54 | 1.2936 | 0.0245 | 1.23 | 0.01 |
| **EC14** | 3219.4 | 158.67 | 10758 | 876.73 | 3.3472 | 0.324 | 51.24 | 1.288 |
| **EC15** | 473.26 | 16.404 | 809.52 | 31.188 | 1.7135 | 0.1277 | 2.789 | 0.01 |
| **EC16** | 4586.3 | 67.924 | 34407 | 4153.8 | 7.7022 | 2.1242 | 0.7261 | 1.988 |
| **EC17** | 1041.6 | 61.392 | 6183.2 | 121.45 | 5.959 | 0.5236 | 7.204 | 0.9196 |

**Table S4. Plasmids used in this study.**

| Plasmid name | Relevant characteristics |
|---|---|
| **pSEVA131** | Cloning vector, oriV (pBBR1), AmR from SEVA collection. |
| **pTBAB+** | Initial coupled biosensor vector bearing $P_{tph}$-tphR pair from *Camomonas sp.* E6; oriV (pBBR1), AmR |
| **pTBAB1** | Initial coupled biosensor vector bearing $P_{tph}$-tphR pair from *P. litoralis* identified from homology analysis; oriV (pBBR1), AmR |
| **pTBAB2** | Initial coupled biosensor vector bearing $P_{tph}$-tphR pair from *T. baoligensis* identified from homology analysis; oriV (pBBR1), AmR |
| **pTBAB3** | Initial coupled biosensor vector bearing $P_{tph}$-tphR pair from *L. curvus* identified from homology analysis; oriV (pBBR1), AmR |
| **pTBAB4** | Initial coupled biosensor vector bearing $P_{tph}$-tphR pair identified from *Z. caldifontis* identified from homology analysis; oriV (pBBR1), AmR |
| **pTB+** | Decoupled biosensor, with $P_{tph}$-tphR pair from *Camomonas sp.* E6; oriV (pBBR1), AmR |
| **pTB4** | Decoupled biosensor, with $P_{tph}$-tphR pair from *Z. caldifontis.* This biosensor was used as template to all subsequent constructs; oriV (pBBR1), AmR |
| **pTB4-Lib** | $P_{out}$ library vector; pTB4 was used as template for introduction of the $P_{out}$ library degenerate oligo GA229; resulting *Lib* mutants spanned 1-5,000 clones. |
| **pDR1-17** | Final TPA biosensor vector; assembled as depicted in Table 1. The $P_{out}$ in these vectors corresponds to those ranked by basal transcriptional strength. oriV (pBBR1), AmR |
| **pEC1-17** | Final TPA biosensor vector; assembled as in depicted Table 1. The $P_{out}$ in these vectors corresponds to those ranked by promoter sensitivity. oriV (pBBR1), AmR |
| **pVEC1** | Final validation biosensor construct, where $RBS_{out}$ was set to the highest level (+1), $P_{reg}$ was set to -0.7 and the level of $P_{out}$ was set to the advised mid-lower point (-0.3), while $RBS_{trans}$ was maintained at -1 |

**Table S5. List of primers and oligos used in this study.**

| Primer name | Sequence |
|---|---|
| **GA6** | CTCGAGGTTTGACAGCTTATC |
| **AB27** | ATGAGCAAAGGTGAAGAACTGTTTAC |
| **GA229** | GCAGTCGATGATAAGCTGTCAAACCTCGAGggcgaggccttgcgttg<br>cagcagAAGTkTkCGhATGdCGMAMATCTagcTNGTNNcgggtcaaaccggcg<br>acNANANTtcaaaggccccTTTAACTTTAAGAAGGGTGTATACATATGAGCA<br>AAGGTGAAGAACTGTTTACCGGT |
| **GA86** | TTTAACTTTAAGAAGGAGGTATACATATGCTCCCGGAGTCCAAG |
| **GA87** | ATGTATACCTCCTTCTTAAAGTTAAATTACAGACCCTGAGGATACAGTTTTT |
| **GA88** | ATGTATACATGTTTCTTAAAGTTAAATTACAGACCCTGAGGATACAGTTTTT |
| **GA89** | TTTAACTTTAAGAAACATGTATACATATGCTCCCGGAGTCCAAG |
| **GA90** | ATGTTTTTCCTCATTATAAAGTTAATCTTACAGACCCTGAGGATACAGTTTTT |
| **GA91** | GATTAACTTTATAATGAGGAAAAACATATGCTCCCGGAGTCCAAG |
| **GA92** | ATGTTTTTCCTGTTTATAAAGTTAATCTTACAGACCCTGAGGATACAGTTTTT |
| **GA93** | GATTAACTTTATAAACAGGAAAAACATATGCTCCCGGAGTCCAAG |
| **GA94** | ATGTTTTTGAACCGTATAAAGTTAATCTTACAGACCCTGAGGATACAGTTTTT |
| **GA95** | GATTAACTTTATACGGTTCAAAAACATATGCTCCCGGAGTCCAAG |
| **GA96** | ATGTTTTTTAACCATATAAAGTTAATCTTACAGACCCTGAGGATACAGTTTTT |
| **GA97** | GATTAACTTTATATGGTTAAAAAACATATGCTCCCGGAGTCCAAG |
| **GA102** | ATGCAGGATAAAAACTTTGTTGAAAGC |
| **GA115** | ATGAGCAAAGGTGAAGAACTGTTTACCGGT |
| **GA116** | GGTTTGACAGCTTATCATCGACTGC |
| **GA201** | caccttgatttgatcgctCTC |
| **GA202** | TTTAACTTTAAGAAGGGTGTATACATATGAGC |
| **GA216** | GCGGCTTGGACTCCGGGAGCATatgTTTTTCCTCCTTATAAAGTTAATCTTAGAGCATGGCGGTC<br>AGC |
| **GA217** | GTTTAACTTTGAAATAAGGAGGTAATACAAATGGTCGAGGAAGTGCG |
| **GA218** | TTGTATTACCTCCTTATTTCAAAGTTAAACAAAAT |
| **GA219** | ggggcctttgaagtgtggtc |
| **GA220** | ATGTATATGACTATCTTAAAGTTAAAggggcctttgaagtgtggtc |
| **GA221** | ATGTATACACCCTTCTTAAAGTTAAAggggcctttgaagtgtggtc |
| **GA222** | ATGTATATCTCCTTCTTAAAGTTAAAggggcctttgaagtgtggtc |
| **GA223** | GAGCTGCAGCGGCTTGGACTCCGGGAGCATtagaaaacctccttagcatgattaagatgtttcagtacgaaa<br>attCCAGCCAGAAATCATCCTTAGCGAAAGCTAAGGATTTTTTTTATCTG |
| **GA224** | GAGCTGCAGCGGCTTGGACTCCGGGAGCATtagaaaacctccttagcatgattaagatgtttcagtacgaaa<br>attTAGATTAATTAAACCAGCCAGAAATCATCCTTAGCGAAAGCTAAGGATTTTTTTTATCTG |
| **GA225** | GAGCTGCAGCGGCTTGGACTCCGGGAGCATtagaaaacctccttagcatgattaagatgtttcagtacgaaa<br>attGCTTAATTAAACCAGCCAGAAATCATCCTTAGCGAAAGCTAAGGATTTTTTTTATCTG |
| **GA226** | TTAGCGAAAGCTAAGGATTTTTTTTATCTGTTAGTCCTGATAGATCTGCTCC |
| **GA227** | TTAGCGAAAGCTAAGGATTTTTTTTATCTGTTAGAGCATGGCGGTCAGC |
| **GA228** | GAGCTGCAGCGGCTTGGACTCCGGGAGCATtagaaaacctccttagcatgattaagatgtttcagtacgaaa<br>attAAATAAATTAATTAAACCAGCCAGAAATCATCCTTAGCGAAAGCTAAGGATTTTTTTTATCTG |
| **GA232** | GCATATGTATACACCCTTCTTAAAGTTAAAggTACATACATTATATCCATGTCAACC |
| **GA233** | TAccTTTAACTTTAAGAAGGGTGTATACATATGCTCCCGGAGTCCAAG |
| **GA234** | GCATATGTATATCTCCTTCTTAAAGTTAAAggTACATACATTATATCCATGTCAACC |
| **GA235** | TAccTTTAACTTTAAGAAGGAGATATACATATGCTCCCGGAGTCCAAG |
| **GA236** | GCATATGTATATGACTATCTTAAAGTTAAAggTACATACATTATATCCATGTCAACC |
| **GA237** | TAccTTTAACTTTAAGATAGTCATATACATATGCTCCCGGAGTCCAAG |
| **PoutFW** | caaaggccccTTTAACTTTAAGA |
| **21C5** | GGCGAGGCCTTGCGTTGCAGCAGAAGTTTGCGAATGTCGAACATCTAGCTTGTCACG |

| | |
|---|---|
| **(DR $P_{out}$ +1)** | GGTCAAACCGGCGACTATAATTCAAAGGCCCC |
| **30A5** | GGCGAGGCCTTGCGTTGCAGCAGAAGTGTGCGAATGTCGAAAATCTAGCTCGTCGCG |
| **(DR $P_{out}$-1 & EC $P_{out}$ +1)** | GGTCAAACCGGCGACTATAATTCAAAGGCCCC |
| **45E7** | GGCGAGGCCTTGCGTTGCAGCAGAAGTGTTCGCATGTCGCAAATCTAGCTTGTGACGG |
| **(EC and DR $P_{out}$ 0)** | GTCAAACCGGCGACTATAGTTCAAAGGCCCC47 |
| **48G6** | GGCGAGGCCTTGCGTTGCAGCAGAAGTTTTCGTATGACGAAAATCTAGCTTGTCACGGG |
| **(EC $P_{out}$ -1)** | TCAAACCGGCGACTAGAGTTCAAAGGCCCC |

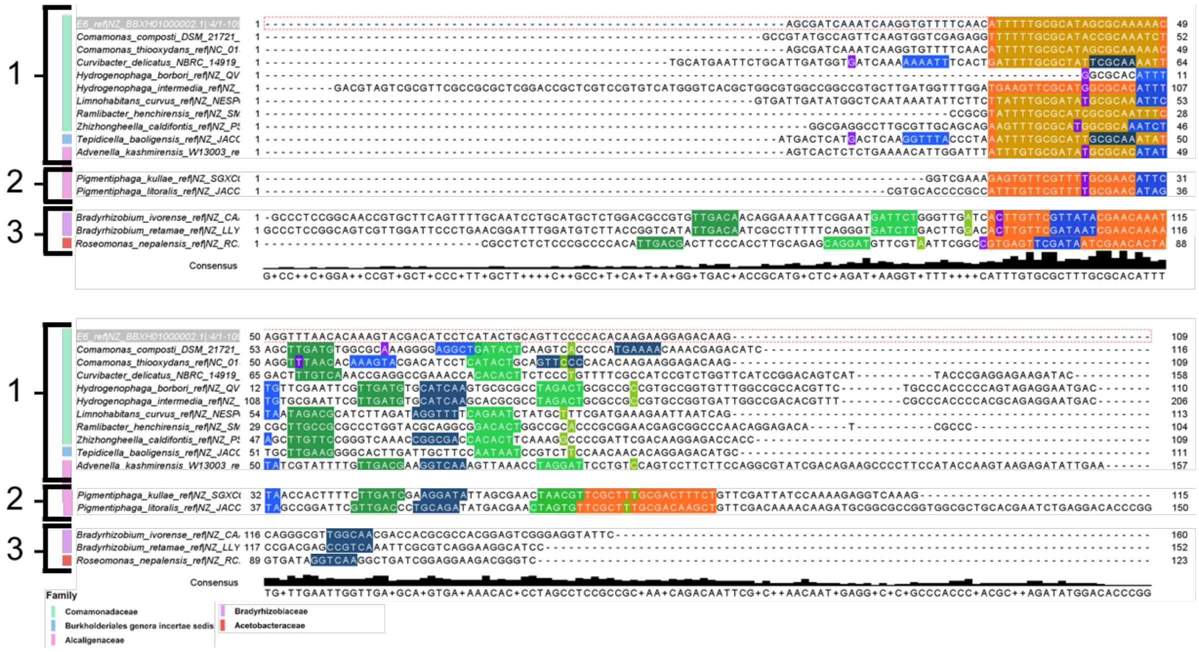

**Figure S1. Distribution of TSSs and hex-boxes inferred using the extended thermodynamic model within *tph* and *tph*-like promoters.** Putative promoter regions from *tph* and *tph*-like operons were aligned using Clustal Omega. Hits to Kasai et al.'s TphR binding site motif are shown as yellow boxes within the sequence. Hits to the MEME generated putative TphR binding site are shown in orange. The predicted location of the hexboxes for this *tphC*-TSS are shown in light green (-10 site) and dark green (-35 site). On the reverse strand (towards *tphR*), the strongest TSS predicted is shown in light purple. The predicted location of the hexboxes for this *tphR*-TSS are shown in light blue (-10 site) and dark blue (-35 site). *Comamonas* strain sp. E6's promoter region is highlighted in grey at the top of the list of promoters, and its sequence is shown surrounded by a dashed horizonal red line. The taxonomic family of the bacteria that contain each of these operons is shown to the left of the promoter's name. The operons are grouped into sets based on their configuration; this is shown to the extreme left of the diagram. Two operons have not been plotted due to a lack of TSS mapping data, and the consensus sequence shown is therefore not totally accurate across the whole promoter region.

**a**

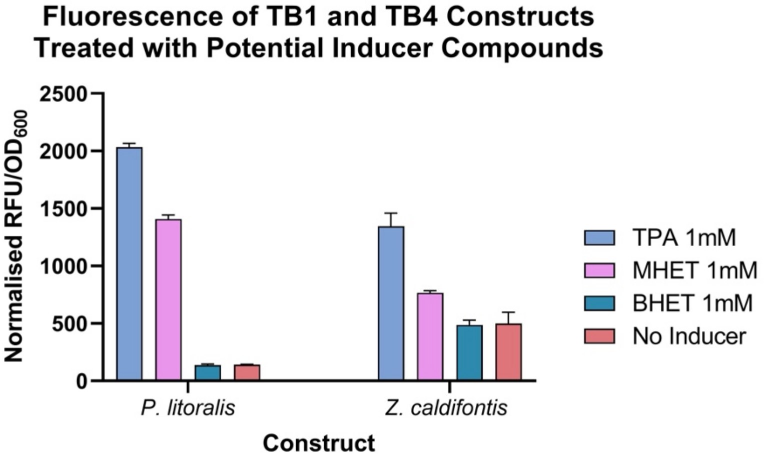


**b**

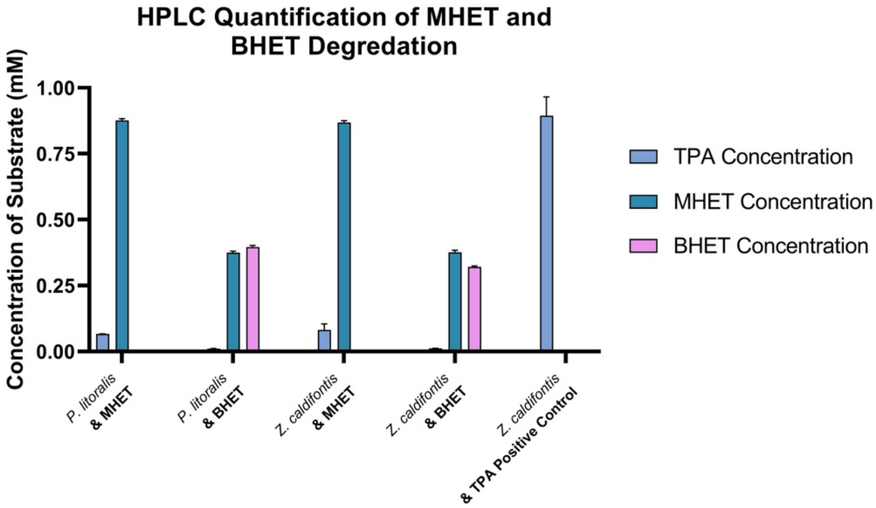


**Figure S2. TPA Biosensors treated with PET breakdown products**. (**a**) TB1 and TB4 biosensors were treated with MHET, BHET or TPA. Induction was seen upon treatment with TPA or MHET, but not with BHET. (**b**) Potential hydrolysis of BHET, MHET into TPA by the *P. putida* host, containing the *P. litoralis* and *Z. caldifontis* biosensors were analysed by HPLC. BHET is roughly 50% converted to MHET after the 16h incubation with slight further conversion to TPA. Similarly, added MHET is partially to TPA after the 16h incubation. In light of the partial conversion of MHET to TPA during incubation and the lack of induction when treated with BHET, we believe the induction upon treatment with MHET to be due to TPA conversion over the induction period. Data is shown as mean and standard deviation of 3 biological replicates for all conditions other than the *Z. caldifontis* + TPA positive control for which n= 2.

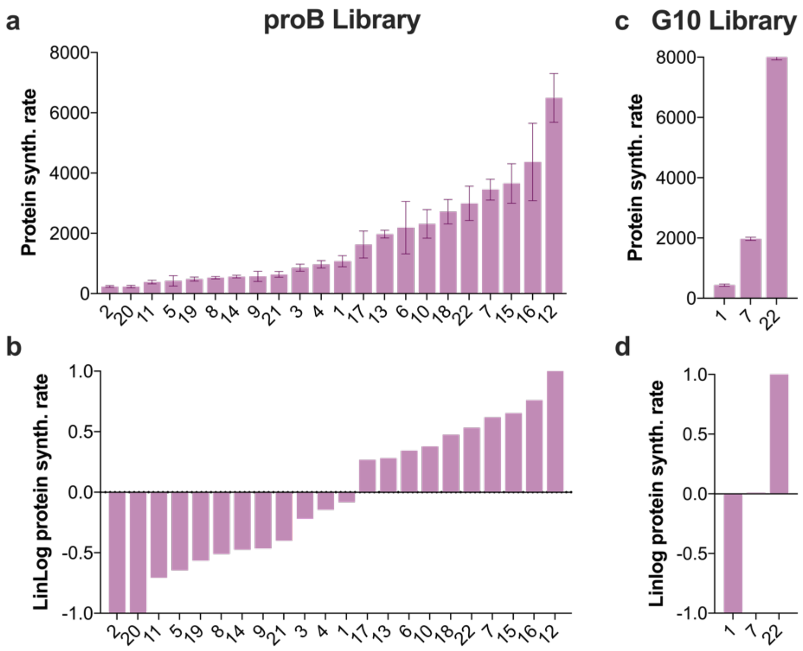


**Figure S3. Constitutive ProB promoter and RBS G10 libraries activity in *P. putida*.** Obtained protein synthesis rates for the constitutive proB (a) and G10 promoter (c) libraries. Synthesis rates from both libraries were transformed into logarithmic scale values (lin log transformation) to establish their ranking order in *P. putida* (b, d). All data are the mean ±SD of three biological triplicates.

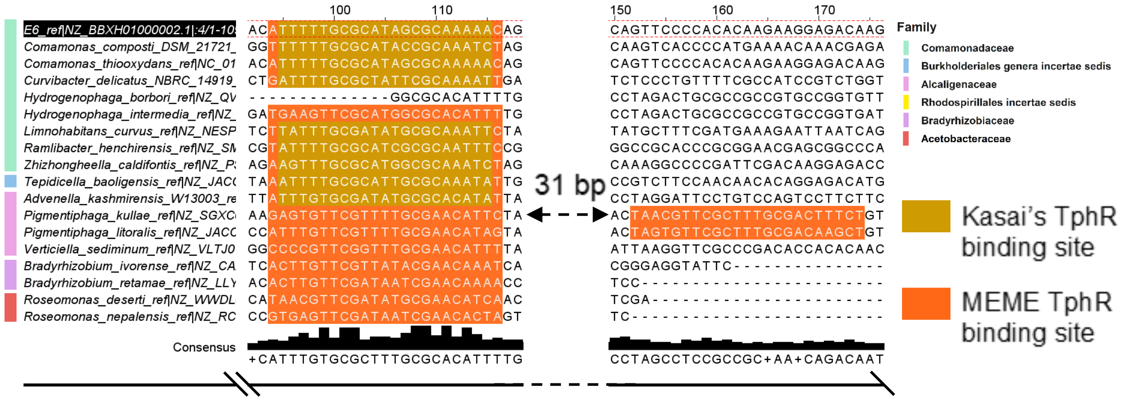


**Figure S4. Distribution of putative TphR binding sites within *tph* and *tph*-like promoters.** Putative promoter regions from *tph* and *tph*-like operons were aligned using benchling. Hits to Kasai et al.'s TphR binding site motif are shown as yellow boxes within the sequence. Hits to the MEME generated putative TphR binding site are shown in orange. *Comamonas* strain sp. E6's promoter region is highlighted in black at the top of the list of promoters, and its sequence is shown surrounded by a dashed horizonal red line. The taxonomic family of the bacteria that contain each of these operons is shown to the left of the figure.

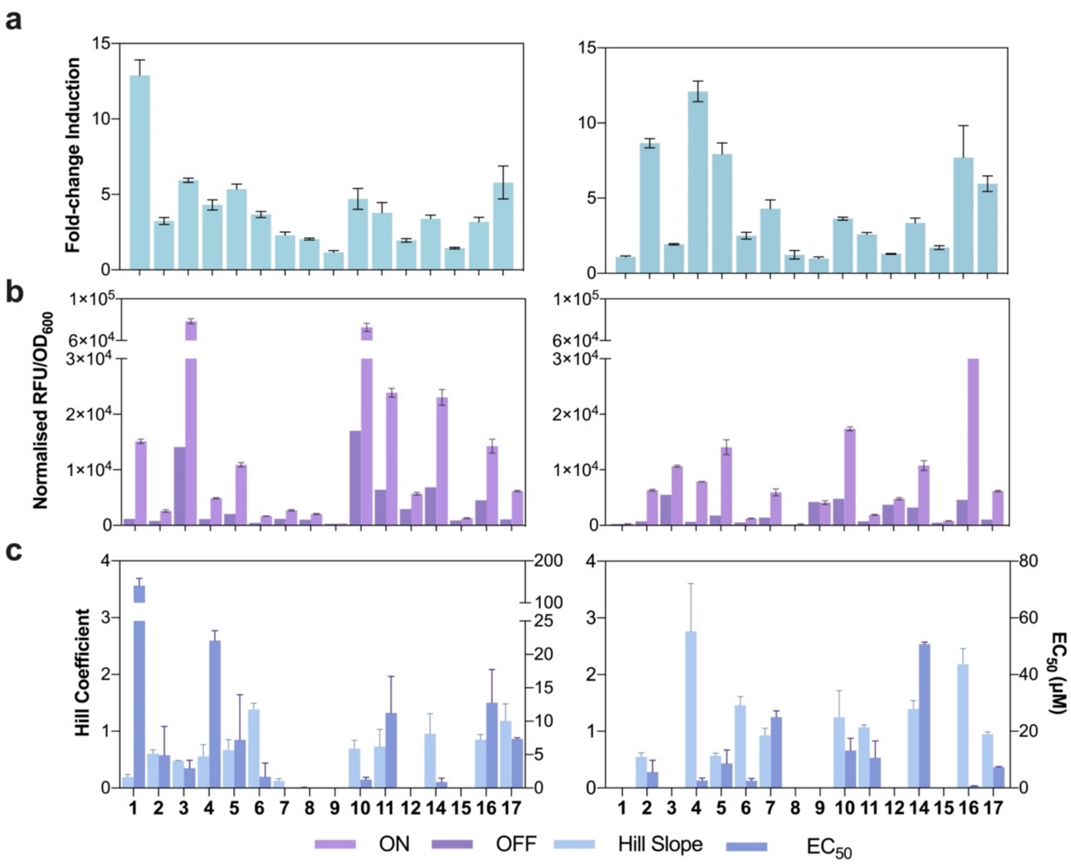


**Figure S5. Full experimental trials data for DR and EC experiments**. (a) Fold-change induction obtained between the maximal ON (1 mM) and OFF (no TPA) for both the DR (left panel) and EC (right panel) ranking experiments. (b) Normalised GFP fluorescence at both the ON and OFF states for both rankings. (c) Hill coefficient and $EC_{50}$ values obtained from titration data of both the DR (left panel) and EC (right panel) rankings. All data shown are the mean ±SD of three biological triplicates.

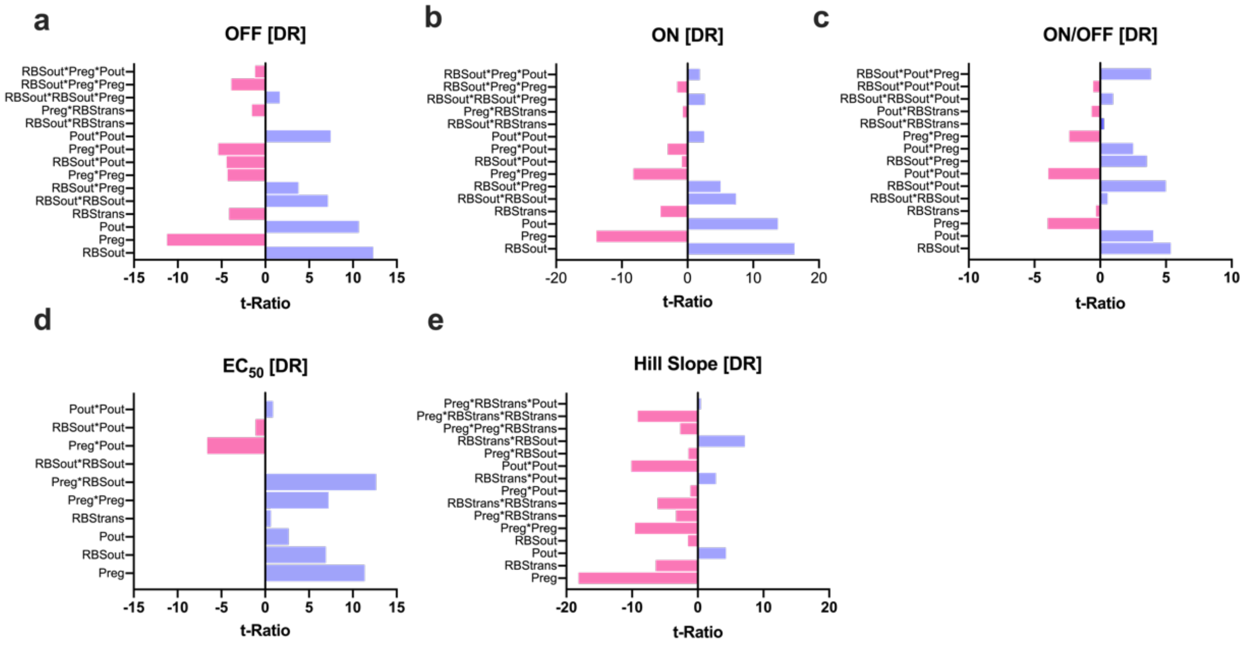


**Figure S6. Factor screening and selection from DR experiment**. (a-e) Lenth t-ratio of each response (a) OFF, (b) ON, (c) ON/OFF, (d) $EC_{50}$ and (e) Hill Slope, showing those factors deemed important by JMP two-level screening analysis for the DR experiment. T-ratios are used to assess factor importance. The colour of the corresponding bars indicates the effect from each factor or interaction upon the corresponding response; where light pink represents negative effect and purple represents a positive effect. Only those factors that were significant at the 0.1 confidence interval were deemed significant and included in the model.

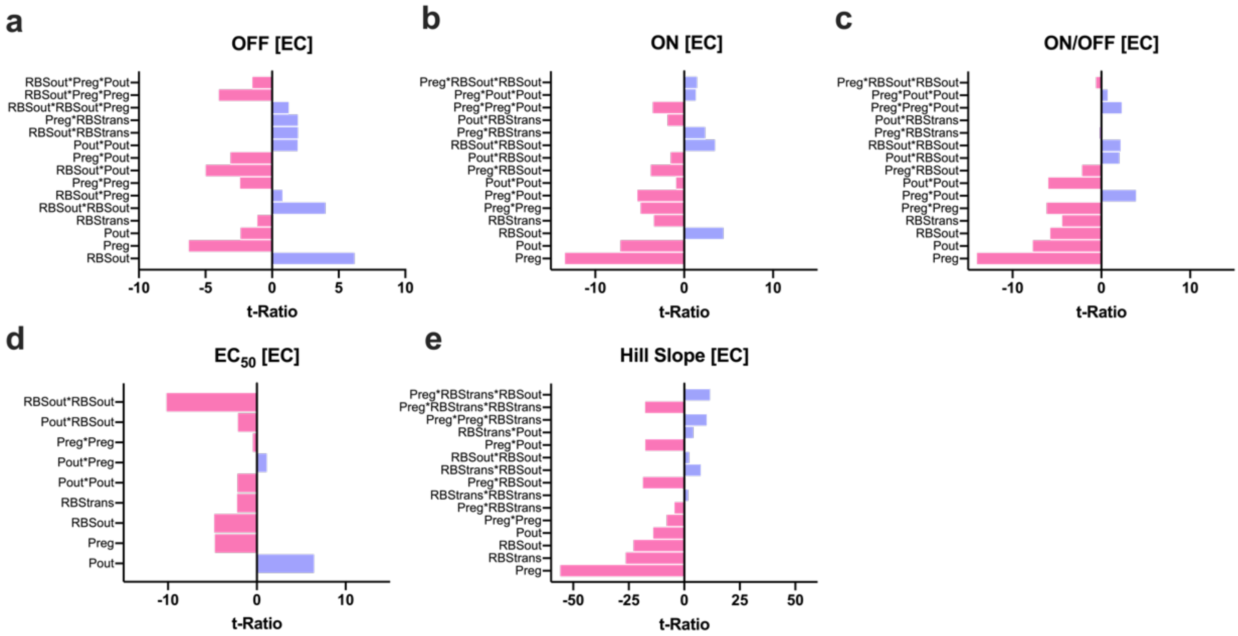


**Figure S7. Factor screening and selection from EC experiment**. (a-e) Lenth t-ratio of each response (a) OFF, (b) ON, (c) ON/OFF, (d) $EC_{50}$ and (e) Hill Slope, showing those factors deemed important by JMP two-level screening analysis for the EC experiment. T-ratios are used to assess factor importance. The colour of the corresponding bars indicates the effect from each factor or interaction upon the corresponding response; where light pink represents negative effect and purple represents a positive effect. Again, only those factors that were significant at the 0.1 confidence interval were deemed significant and included in the model.

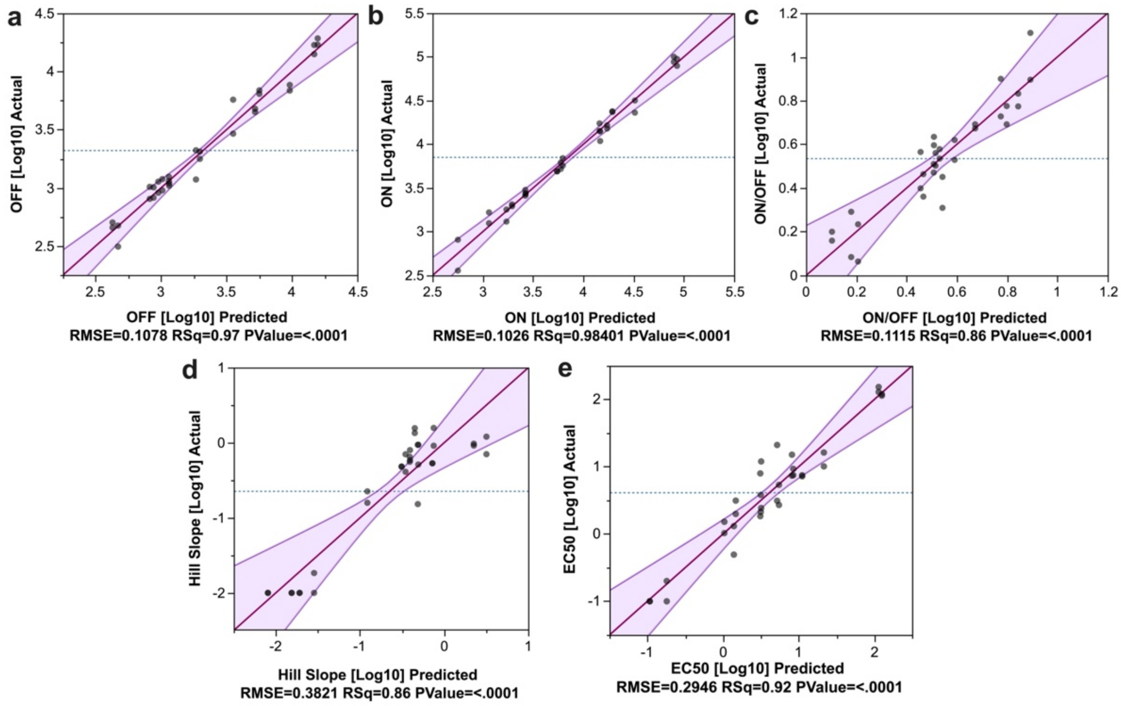


**Figure S8. Standard Least Squares Regression model performance for the DR experiment**. Actual versus predicted scatter plots showing the performance of the SLSR when predicting (a) OFF, (b) ON, (c) ON/OFF, (d) Hill Slope and (e) $EC_{50}$ of the DR DSD experiment. The model shows good prediction for all responses (all cases p <0.0001; $R^2$ > 0.86).

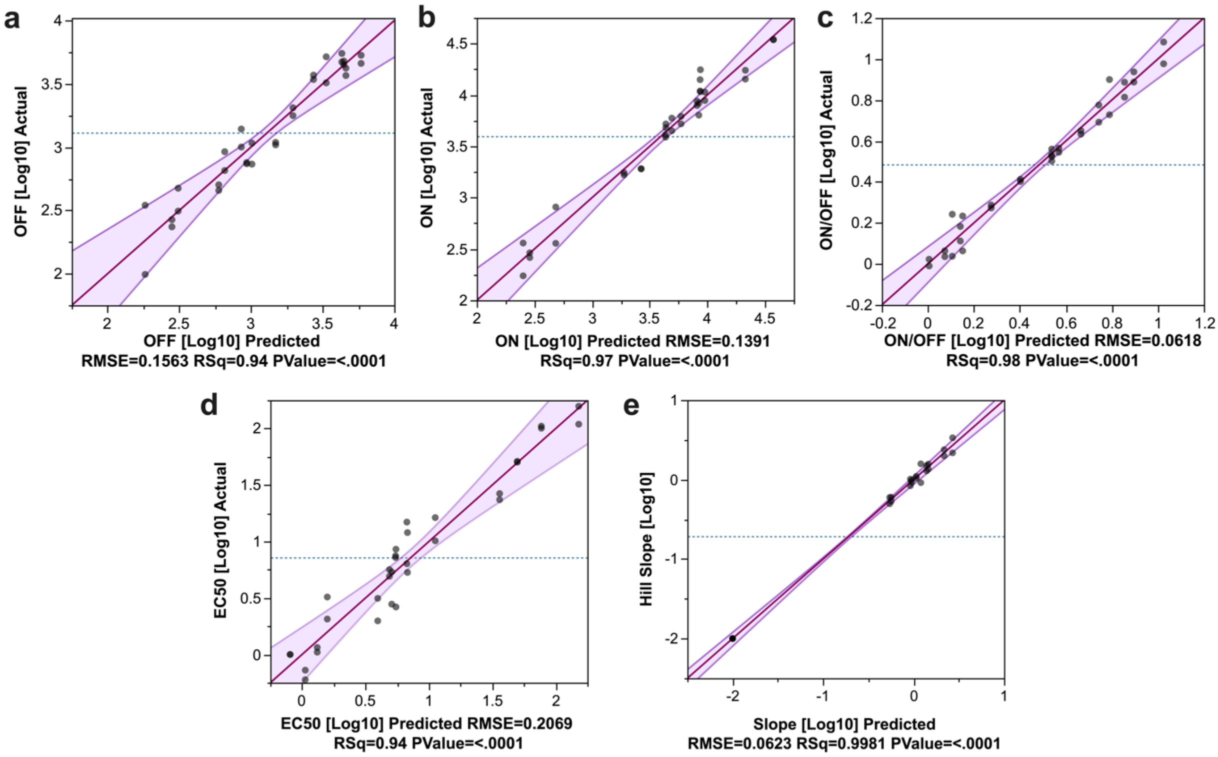


**Figure S9. Standard Least Squares Regression model performance for the EC experiment**. Actual versus predicted scatter plots showing the performance of the SLSR when predicting (a) OFF, (b) ON, (c) ON/OFF, (d) Hill Slope and (e) $EC_{50}$ of the DR DSD experiment. The model shows good prediction for all responses (all cases p <0.0001; $R^2$ > 0.86).

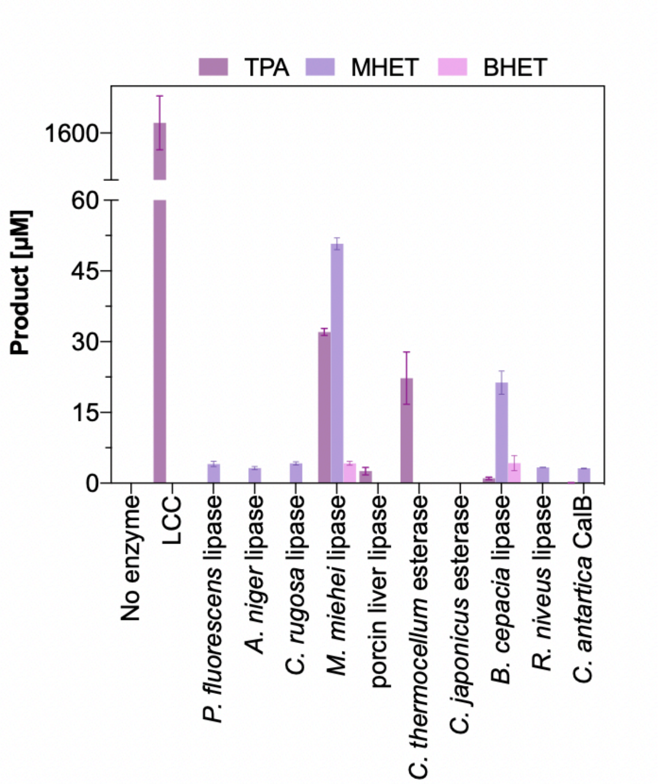


**Figure S10. Esterase and lipase activity towards PET film**. HPLC quantification results of TPA, MHET and BHET products from enzyme screening was assayed using 10 % (w/v) solid loading in 100 mM sodium phosphate buffer pH 7.0 with 25 mg enzyme/g PET. Assays were incubated at 40 °C and 125 rpm for 72 hours before being filtered for HPLC analysis.

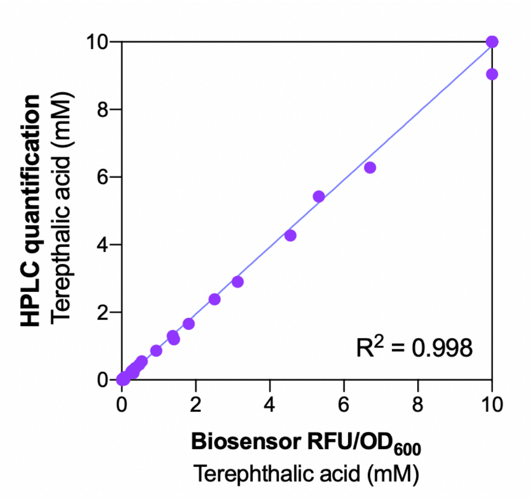


**Figure S11. Biosensor- versus HPLC-mediated quantification of TPA.** Both methodologies showed excellent correlation ($R^2$ = 0.998) of TPA quantification from PET hydrolysates in the enzyme condition test. The DR1 calibration curve is able to accurately determine the concentration of released TPA as compared to HPLC,
